# Structural Basis of MicroRNA Biogenesis by Dicer-1 and Its Partner Protein Loqs-PB

**DOI:** 10.1101/2022.04.19.488762

**Authors:** Karina Jouravleva, Dmitrij Golovenko, Gabriel Demo, Robert C. Dutcher, Traci M. Tanaka Hall, Phillip D. Zamore, Andrei A. Korostelev

## Abstract

In animals and plants, Dicer enzymes collaborate with double-stranded RNA-binding proteins to convert precursor-microRNAs (pre-miRNAs) into miRNA duplexes. We report six cryo-EM structures of *Drosophila* Dicer-1 and its partner Loqs-PB. The structures show Dicer-1•Loqs-PB (1) before binding pre-miRNA, (2) after binding and in a catalytically competent state, (3) after nicking one arm of the pre-miRNA, (4) following complete dicing and initial product release. Our reconstructions suggest that pre-miRNA binds a rare, open conformation of the Dicer-1•Loqs-PB heterodimer, enabling conformational proofreading. The Dicer-1 dsRBD and three Loqs-PB dsRBD domains form a tight belt around the pre-miRNA, distorting the RNA helix to place the scissile phosphodiester bonds in the RNase III active sites. Pre-mRNA cleavage shifts the RNA-binding domains and tightens Dicer-1, promoting product release. Our data suggest a model for how the Dicer-1•Loqs-PB complex effects a complete cycle of pre-miRNA recognition, stepwise endonuclease cleavage, and sequential product release.

## INTRODUCTION

In plants and animals, microRNAs (miRNAs) direct Argonaute (AGO) proteins to repress mRNA expression by inhibiting translation or marking the RNA for destruction (Chen, 2009; Bartel, 2018; Dong et al., 2022; Zhang et al., 2022). In animals, miRNA biogenesis begins in the nucleus with the excision of a pre-miRNA stem loop from a primary miRNA transcript by the ribonuclease III enzyme Drosha, aided by a partner protein with multiple double-stranded RNA-binding domains (dsRBDs) (Lee et al., 2003; Denli et al., 2004; Gregory et al., 2004; Han et al., 2004; Han et al., 2006). The pre-miRNA is then exported to the cytoplasm, where a second ribonuclease III enzyme, Dicer, in complex with its dsRBD-containing partner protein Loquacious in flies and TRBP or PACT in mammals, converts the pre-miRNA to a duplex comprising a miRNA guide paired to its miRNA* passenger strand (Grishok et al., 2001; Hutvágner et al., 2001; Knight and Bass, 2001; Chendrimada et al., 2005; Förstemann et al., 2005; Haase et al., 2005; Maniataki and Mourelatos, 2005; Saito et al., 2005; Lee et al., 2006). The miRNA duplex is loaded into an AGO protein, and the miRNA*—the strand of the duplex equivalent to the target RNA—is evicted from the complex (Khvorova et al., 2003; Schwarz et al., 2003; Meister et al., 2004; Okamura et al., 2004; Förstemann et al., 2007; Tomari et al., 2007; Kawamata et al., 2009). Because miRNA-guided AGO proteins find their targets via base pairing between an mRNA and a small region of sequence at the miRNA 5′ end, the seed (nucleotides g2–g8), even a single nucleotide shift in the miRNA 5′ end alters the entire repertoire of miRNA targets (Lai, 2002; Lewis et al., 2003; Brennecke et al., 2005; Chiang et al., 2010; Fukunaga et al., 2012). Thus, accurate pre-miRNA processing by Dicer is a prerequisite for miRNA function.

Dicer comprises head, core, and base superdomains (Lau et al., 2009; Wang et al., 2009; Lau et al., 2012; Taylor et al., 2013; Liu et al., 2018; Sinha et al., 2018; Wang et al., 2021; Wei et al., 2021). The head contains a PAZ domain, which binds the two-nucleotide, 3′ overhang characteristic of pre-miRNAs (Lingel et al., 2003; Song et al., 2003; Yan et al., 2003; Lingel et al., 2004; Ma et al., 2004). The pre-miRNA is cleaved by the intramolecular RNase III dimer that forms the lower half of the Dicer core (Macrae et al., 2006; MacRae et al., 2007; Park et al., 2011). The C-terminal dsRBD in the Dicer core is proposed to bind the dsRNA substrate, allowing it to engage the intramolecular RNase III dimer (Nicholson, 2014; Hansen et al., 2019). A catalytically inactive DExD/H-box helicase domain forms the Dicer base, which recognizes the pre-miRNA loop and provides a platform to bind Dicer partner proteins (Lee et al., 2006; Daniels et al., 2009; Cenik et al., 2011; Tsutsumi et al., 2011; Ota et al., 2013). An αβββα structural motif (domain of unknown function [DUF] 283) in the base resembles a canonical dsRBD (Gleghorn and Maquat, 2014), but binds single-stranded RNA and Dicer partner proteins, not dsRNA (Qin et al., 2010; Ota et al., 2013; Kurzynska-Kokorniak et al., 2016).

In *Drosophila melanogaster*, Dcr-1, the ortholog of mammalian Dicer, collaborates with the dsRBD protein Loquacious to process pre-miRNAs into miRNAs (Förstemann et al., 2005; Jiang et al., 2005; Saito et al., 2005). Alternative splicing generates three *loqs* isoforms that encode Loqs-PA, PB, and PD. Dcr-1 relies on Loqs-PA and Loqs-PB for specific, efficient processing of pre-miRNAs (Förstemann et al., 2005; Jiang et al., 2005; Saito et al., 2005; Fareh et al., 2016). Loqs-PB enhances dicing of pre-miRNAs containing mismatches near the scissile phosphates (Tsutsumi et al., 2011). Such mismatches promote loading of the miRNA/miRNA* duplex into AGO for miRNAs residing on the 3′ arm of the pre-miRNA stem (Khvorova et al., 2003; Schwarz et al., 2003). Consequently, Loqs-PB allows Dicer’s preference for perfect stems to co-exist with the thermodynamic asymmetry required for the 3′ pre-miRNA arm to produce miRNAs. Loqs-PB comprises three canonical αβββα dsRBDs: the first two bind dsRNA while the third binds Dcr-1 (Figure 1A) (Haase et al., 2005; Förstemann et al., 2007; Ye et al., 2007; Daniels et al., 2009; Yang et al., 2010; Wilson et al., 2015; Jakob et al., 2016).

**Figure 1.**
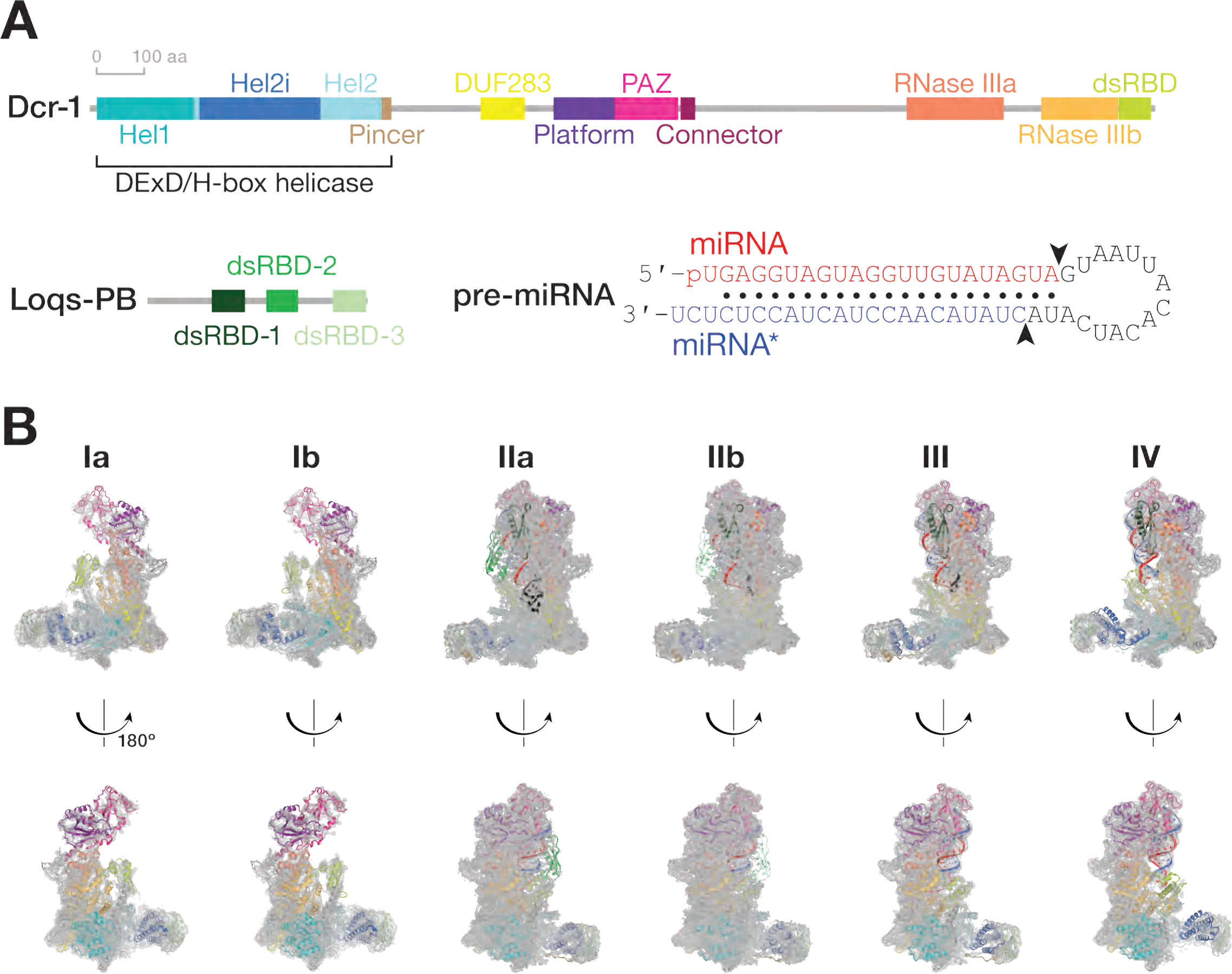
Cryo-EM captures Dcr-1•Loqs-PB through the cycle of processing. **(A)** Domain architecture of *D. melanogaster* Dcr-1 and Loqs-PB proteins and the sequence of the model pre-miRNA used for structure determination. **(B)** Cryo-EM density maps (gray mesh) and the superimposed models for six states of Dcr-1•Loqs-PB alone or in complex with pre-miRNA. Domains are colored as in **(A)**. See also Figures S1–S3.

Structures of *Giardia intestinalis* Dicer, *Drosophila* Dicer-2, and plant and human Dicers alone or bound to dsRNA or pre-miRNA in inhibited states—e.g., by substituting Ca^2+^ for Mg^2+^—reveal the protein’s overall architecture and how this protein family may bind its substrates (Macrae et al., 2006; Liu et al., 2018; Sinha et al., 2018; Wang et al., 2021; Wei et al., 2021). However, it remains unknown how Dicer proteins collaborate with their dsRNA-binding partner proteins to authenticate, bind, and cleave pre-miRNA. We used cryo-EM to visualize how *Drosophila* Dcr-1 collaborates with Loqs-PB to process pre-miRNAs. Here, we describe six structures of the Dcr-1•Loqs-PB complex alone or bound to a model pre-miRNA, at average resolutions from 3.0 to 4.0 Å (Figures 1 and S1–S3; Table S1). These structures show the stepwise mechanism of dicing, from initial substrate binding and positioning of the pre-miRNA in the two RNase III catalytic sites, to the sequential hydrolysis of two phosphodiester bonds in the pre-miRNA stem and the resulting ordered release of the diced products.

## RESULTS

### Ensemble Cryo-EM Captures Dcr-1•Loqs-PB through the Cycle of Pre-miRNA Processing

We purified full-length Dcr-1•Loqs-PB complex from Sf9 cells and collected three cryo-EM data sets to visualize the complete cycle of pre-miRNA processing. Structures Ia and Ib correspond to ∼4.0 Å cryo-EM reconstructions of the Dcr-1•Loqs-PB complex without RNA (Figures 1 and S1–S3). Cryo-EM of the Dcr-1•Loqs-PB heterodimer with a model pre-miRNA whose miRNA resides on the 5′ arm yielded structure IIa at 3.0-Å resolution. This data set was collected in the presence of Ca^2+^ to preserve protein-RNA interactions but inhibit dicing (Provost, 2002). Structure IIa shows the 60-nt long pre-miRNA A-form dsRNA stem and single-stranded loop bound to Dcr-1 and all three dsRBD domains of Loqs-PB. The staggered pre-miRNA cleavage sites are aligned with the two RNase III active centers, indicating that Dcr-1•Loqs-PB is poised in a cleavage-competent, pre-dicing state. Finally, cryo-EM classification of the heterodimer incubated for 5 min with pre-miRNA in the presence of Mg^2+^ at 25°C yielded an ensemble of structures that could be resolved into three distinct states, structures IIb, III, and IV. Structure IIb, a 3.3-Å reconstruction, is similar to structure IIa and corresponds to a cleavage-competent state (Figure 1B). Structure III, also at 3.3-Å resolution, lacks density for the 5′ arm of the pre-miRNA stem beyond the catalytic center of the Dcr-1 RNase IIIb domain but shows continuous density for the 3′ arm of the stem adjacent to the RNase IIIa domain. Structure III therefore corresponds to a nicked reaction intermediate in which the miRNA 3′ end has been generated but the miRNA* 5′ end has not yet been produced (Figure 1B). Structure IV, a 4-Å reconstruction, lacks RNA density for both arms of the stem beyond the scissile phosphates. Structure IV represents Dcr-1•Loqs-PB after release of the cleaved-off loop but before release of the mature miRNA/miRNA* product (Figure 1B). Collectively, these structures illustrate the stepwise mechanism of pre-miRNA processing and suggest how Loqs-PB assists Dcr-1 to accurately and efficiently generate miRNA/miRNA* duplexes ready to load into AGO proteins.

### Without Pre-miRNA, Dcr-1•Loqs-PB Adopts Closed Conformations

Structures Ia and Ib, the RNA-free Dcr-1•Loqs-PB heterodimer, display the previously defined L-shaped Dicer architecture (Figures 2A and S4A), and multiple domains of the complex share the same general topology found in *Drosophila* Dicer-2 and mammalian and plant Dicers (Macrae et al., 2006; Liu et al., 2018; Sinha et al., 2018; Wang et al., 2021). The core and base of Dcr-1 are well defined, but the lower resolution of the head—comprising the PAZ and platform domains—suggests that this region is dynamic without the pre-miRNA. Like the third dsRBD of TRBP—one of two Loqs orthologs in mammals (Daniels et al., 2009; Wilson et al., 2015)—the Loqs-PB dsRBD-3 binds to Hel2i of the Dcr-1 helicase domain. DsRBD-1 and dsRBD-2 of Loqs-PB are not resolved and are likely mobile. Structures Ia and Ib differ by a ∼10 Å movement of the Dcr-1 C-terminal dsRBD, which lies in different positions at the interface between the core and base in the two structures (Figure S4B). In addition, the head is shifted ∼3 Å closer to the core in structure Ia compared to Ib. Additional lower resolution classes in this data set reveal similar Dcr-1 conformations, with its dsRBD poorly resolved in the vicinity of the positions seen in structures Ia and Ib.

**Figure 2.**
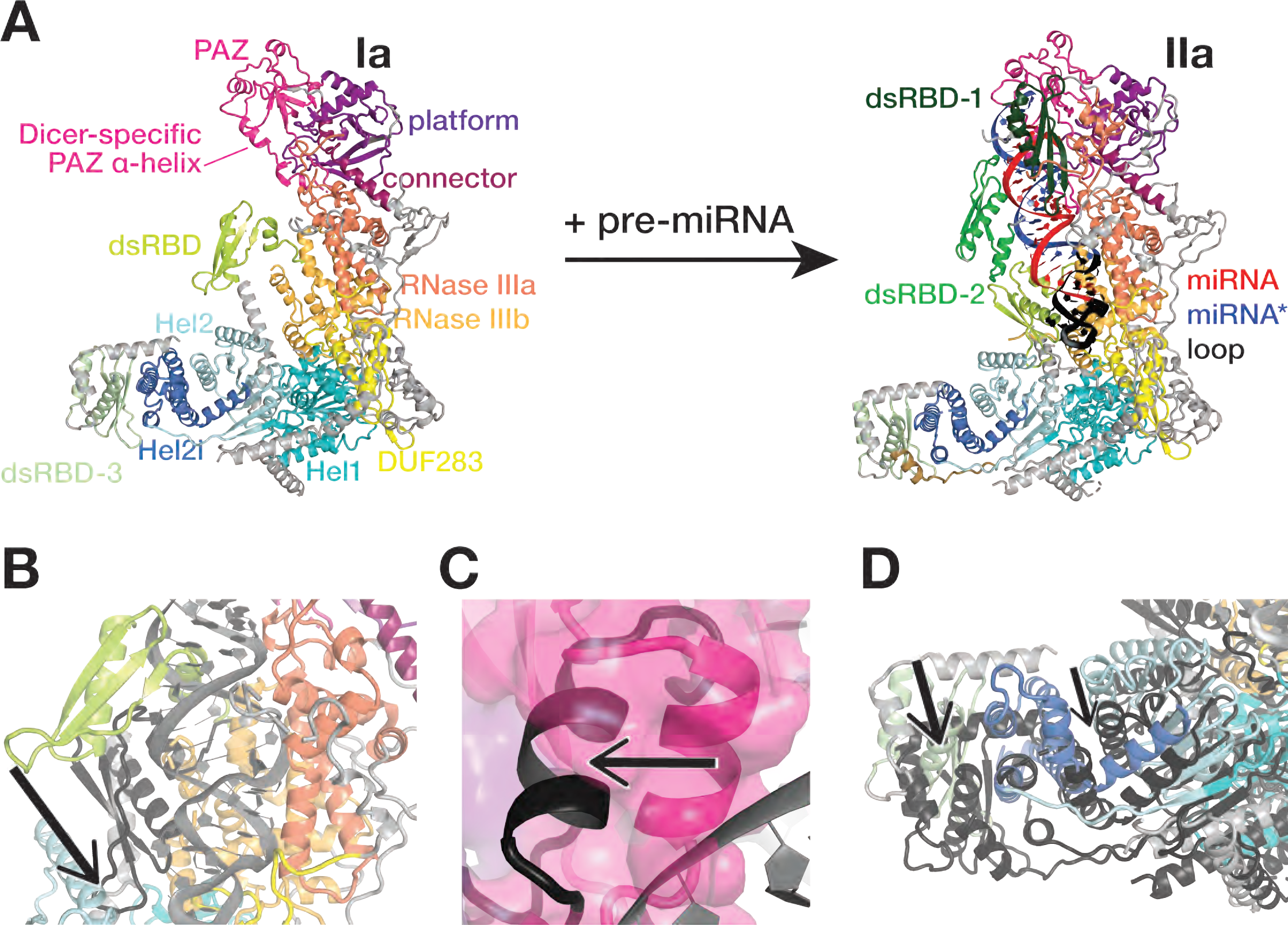
Dcr-1•Loqs-PB conformational equilibrium strongly favors closed states. **(A)** Overall views of the RNA-free Dcr-1•Loqs-PB (structure Ia, gray) and pre-miRNA- bound Dcr-1•Loqs-PB in presence of Ca^2+^ (structure IIa). **(B–D)** Close-up views showing differences in the positions of the Dcr-1 dsRBD **(B)**, the Dicer-specific α-helix of the PAZ domain **(C)**, and the Dcr-1 Hel2i and the Loqs-PB dsRBD-3 domains **(D)** between the RNA-free structure Ia (colored) and the RNA-bound structure IIa (protein domains in dark gray and pre-miRNA in gray). Structural alignments were performed by superposition of Dcr-1•Loqs-PB complexes. Black arrows highlight conformational changes upon pre-miRNA binding. See also Figure S4.

Superposition of the atomic models of RNA-free Dcr-1•Loqs-PB (structure Ia) and RNA-bound Dcr-1•Loqs-PB in a catalytically competent conformation (structures IIa and IIb) reveals three major features of the apo heterodimer. First, the position of the Dcr-1 dsRBD blocks access to the RNase III active sites (Figure 2A). Second, the position of the head differs among structure Ia, structure Ib, and the RNA-bound state (Figures 2A, 2B and S4B), providing further evidence that the head is dynamic before binding of the pre-miRNA termini to the platform and PAZ domains. In both apo structures, the Dicer- specific α-helix of the PAZ domain (aa 1222–1232) packs into the RNA-binding pocket, a configuration incompatible with pre-miRNA binding (Figure 2C). Third, Hel2i in the Dcr-1 base and dsRBD-3 of Loqs-PB lie >10 Å nearer the PAZ domain in the empty heterodimer compared with the RNA-bound complex (Figure 2D). These features suggest that without RNA, the Dcr-1•Loqs-PB conformational equilibrium strongly favors closed states. Accommodation of pre-miRNA requires (1) repositioning the Dcr-1 dsRBD; (2) rearranging and stabilizing the head so that it can bind the 5′ and 3′ ends of the pre-miRNA; and (3) moving Hel2i ≥10 Å to widen the cleft between the core and base to accommodate the pre-miRNA loop.

### The Dcr-1•Loqs-PB Heterodimer Recognizes All of the Pre-miRNA Hallmark Features

Structures IIa and IIb reveal how the Dcr-1•Loqs-PB heterodimer recognizes pre-miRNA. The defining features of a pre-miRNA are a two-nucleotide, single-stranded, 3′ overhanging end, a 5′-monophosphate, and a ∼22 bp double-stranded stem whose arms are linked by a single-stranded loop (Ambros et al., 2003; Park et al., 2011; Bartel, 2018). Consistent with previous biochemical and structural data (Macrae et al., 2006; Tsutsumi et al., 2011; Tian et al., 2014; Liu et al., 2018), the pre-miRNA 3′ overhang nestles in the PAZ domain (Figures 3A and S5A). To accommodate the pre-miRNA, the Dicer-specific α-helix (aa 1222–1232) of the PAZ domain is shifted by ∼15 Å from its position in the RNA-free structure (Figure 2C). The RNA-bound conformation of Dcr-1•Loqs-PB differs from the structures of dicing-incompetent human Dicer (Liu et al., 2018) and murine-specific, ΔHel1 DICER^O^ (Zapletal et al., 2022), in which the 5′ terminal phosphate and first nucleobase are disengaged from the PAZ domain. In structures IIa and IIb, the uracil of the 5′ terminal nucleotide U1 is buried in a binding pocket formed by the PAZ and platform domains, with the nucleobase stacked between Arg994 and Arg1027 (Figure 3B). The π-Arg stacking and lack of steric hindrance at the Watson-Crick face would allow binding of other 5′ nucleotides, in keeping with the occurrence of all four bases as the 5’ nucleotide of *D. melanogaster* and other Dipteran pre-miRNAs. The position of U1 differs from those in isolated PAZ:RNA crystal structures (Tian et al., 2014), where His982 (His1196 in *D. melanogaster*) appears to block the entry of the nucleobase into the binding pocket (Figure S5A). These PAZ structures may reflect interactions sampled transiently during RNA recognition. The pre- miRNA 5′ phosphate is held in a pocket formed by the side chains of Arg1200 and Arg1207 and by the backbone amide groups of Ala1199 and Arg1200. This recognition mechanism contrasts that of plant DCL3, in which the phosphate binds a wider pocket formed by lysine side chains (Figure S5B and S5C) (Wang et al., 2021; Chen et al., 2022).

**Figure 3.**
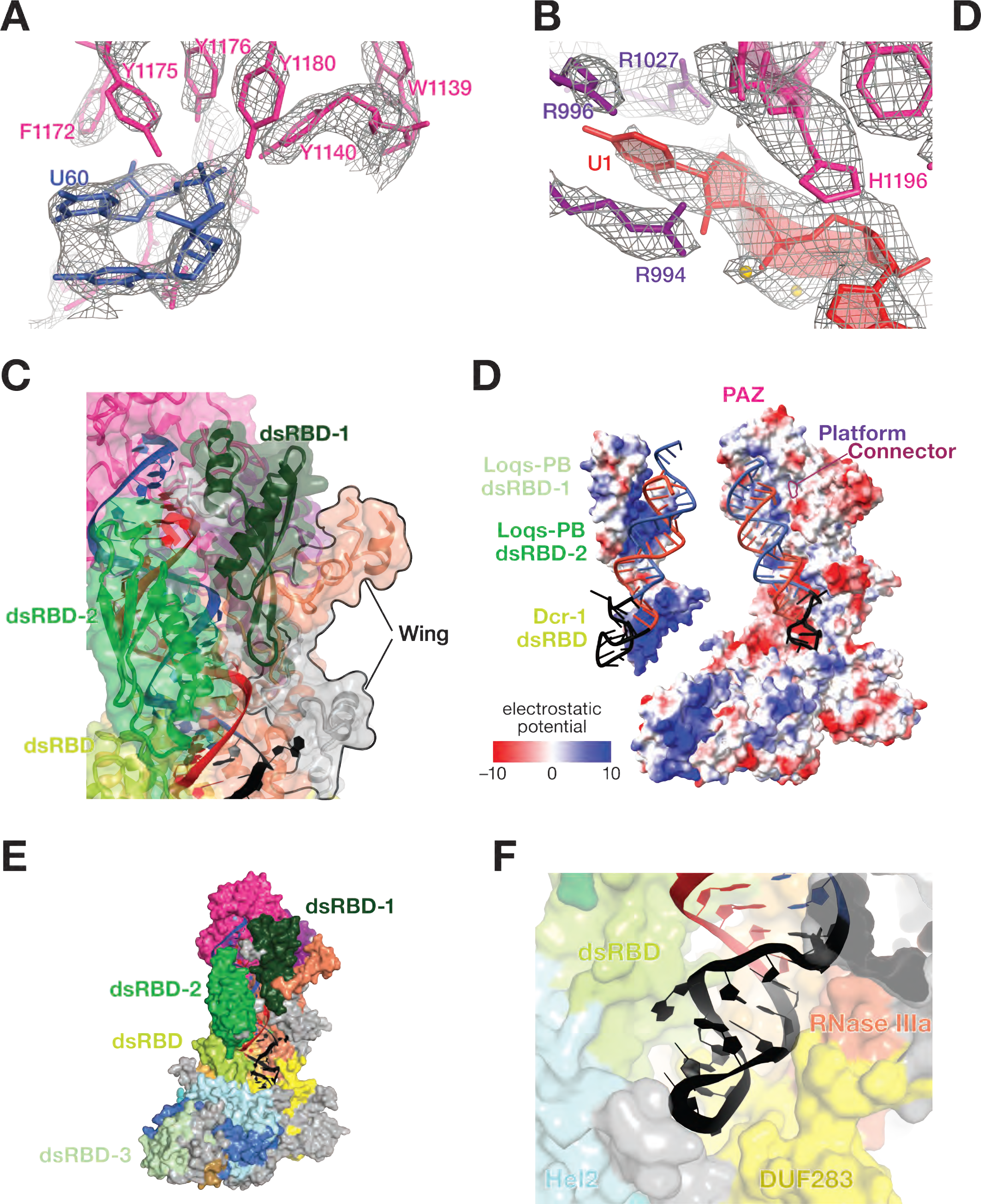
The Dcr-1•Loqs-PB Heterodimer Recognizes All Four Structural Features of Pre-miRNAs. **(A** and **B)** Cryo-EM density (mesh) of the PAZ domain pocket that binds the 3′ overhang of the pre-miRNA **(A)** and the binding pocket formed by the PAZ and platform domains that binds the 5′ terminal nucleotide **(B)**. Ca^2+^ ions are shown as gold spheres. **(C)** Atomic model of Dcr-1•Loqs-PB binding the stem of the pre-miRNA. **(D)** Electrostatic surface view of Dcr-1•Loqs-PB; the RNA is bound in a positively charged channel. Blue, positive charge; red, negative charge. **(E)** Atomic model of Dcr-1•Loqs-PB binding the single-stranded loop of the pre-miRNA. **(F)** Structure of Dcr-1•Loqs-PB•pre-miRNA complex showing multiple domains of Dcr-1 and Loqs-PB enveloping the pre-miRNA. Protein in surface view; RNA in cartoon view All panels correspond to structure IIa. See also Figure S5.

Dcr-1 recognizes the characteristic two-nucleotide, 3′ overhang of the pre-miRNA (C59-U60) using a pocket formed primarily by aromatic residues from the PAZ domain (Figure 3A). Here, the hydroxyl groups of four tyrosine side chains (Tyr1140, Tyr1175, Tyr1176, and Tyr1180) surround the U60 phosphate group, while the β-strand at Ile1236 buttresses the U60 ribose. Moreover, PAZ and pre-miRNA interact with dsRBD-1 of Loqs-PB (Figure 3C), which likely contributes to substrate recognition and stabilization. Helix α1 of Loqs-PB dsRBD-1 (aa 135–145) packs against the minor groove of pre-miRNA helix at G4 and U58, as if tightening the pocket that binds the pre-miRNA termini. At the solvent interface, the Loqs-PB dsRBD-1 is aided by a previously unassigned subdomain of RNase III, which we term the “wing” (aa 924–957 and 1815– 1890; Figure 3C).

The phosphate backbone of the pre-miRNA stem interacts with the positively charged protein extension formed by the platform, PAZ, and connector domains until it reaches the negatively charged catalytic center of Dcr-1 (Figure 3D). On the solvent-exposed side, pre-miRNA is bound to dsRBD-2 of Loqs-PB and the dsRBD of Dcr-1 (Figure 3C). Their positively charged surfaces envelop the pre-miRNA (Figure 3D), as if forming a tight belt that helps to position pre-miRNA in the twin catalytic centers formed by the Dcr-1 intramolecular RNase III dimer (Figure 3E). Binding of the pre-miRNA by tandem dsRBDs—the Dcr-1 dsRBD and dsRBD-2 of Loqs-PB—mirrors the function of the two C-terminal dsRBD domains of *Arabidopsis* DCL3, which produces 24 nt siRNAs from dsRNA (Figures S5B and S5C) (Wang et al., 2021). The Dcr-1 dsRBD is shifted by 10 Å from its position in the RNA-free complex and is wedged between the Dcr-1 core and base, stabilizing the pre-miRNA-bound, open conformation (Figures 2A and 2B).

Finally, Loqs-PB dsRBD-3 binds to Hel2i of the Dcr-1 helicase domain and fastens the Loqs-PB belt around the substrate RNA (Figures 2A and 3F), thus stabilizing and positioning pre-miRNA for cleavage.

The pre-miRNA single-stranded loop (A25–A33) reposes in a pocket formed by the N-terminus and core, base, helicase, DUF283, RNase IIIa and the dsRBD domains of Dcr-1 (Figure 3F). The Hel1 region of the Dcr-1 helicase domain does not directly interact with the pre-miRNA loop, but does support the arrangement of Hel2, DUF283, and RNase IIIa domains. This critical role for Hel1 may explain why its deletion in mouse Dicer leads to widespread somatic changes in the abundance and sequence of miRNAs and increased processing of non-canonical pre-miRNAs with long stems or loops (Zapletal et al., 2022). The cryo-EM structure of murine oocyte-specific Dicer^O^, which lacks Hel1, shows poor density for the helicase and DUF283 domains (Zapletal et al., 2022), perhaps because mouse Hel1 similarly stabilizes the arrangement of the mouse Hel2, DUF283, and RNase IIIa domains.

While the overall shape of the pre-miRNA loop is visible in our cryo-EM density, unambiguous assignment of nucleotide conformations is difficult. The lower resolution of the loop versus the stem is likely due to conformational flexibility, reflecting the ability of Dcr-1 to bind loops of varying sequences and structures. The loop lies in-between the Dcr-1 DUF283 (at Arg837 and Pro837) and the Dcr-1 dsRBD (at Pro2196, Tyr2224 and Arg225). These protein domains appear to provide stacking and electrostatic support for loop nucleotides without necessitating base specificity (Figure 3E). Together, our data show that the Dcr-1•Loqs-PB heterodimer recognizes all four structural features of pre-miRNAs, and suggest a mechanism for how Dcr-1 discriminates between authentic substrates and other hairpins.

We propose that binding the pre-miRNA stabilizes the open—i.e., catalytically competent—conformation of the Dcr-1•Loqs heterodimer, which our data suggests is poorly populated in the absence of RNA. The requirement for pre-miRNA binding to shift the equilibrium from the closed to the open states suggests that Dcr-1•Loqs-PB uses conformational proofreading (Savir and Tlusty, 2007) to authenticate its substrate. Our data suggest that the open conformation is more stable than the closed, catalytically incompetent state only when bound by pre-miRNA and explain how the Dcr-1•Loqs-PB heterodimer rejects inauthentic stem-loop RNAs: such near-cognate substrates bind too weakly to shift the conformational equilibrium from a closed to an open state.

### Dcr-1 and Loqs-PB Position the Pre-miRNA in the Catalytic Center

As in other eukaryotic RNase III enzymes, including human Dicer, human Drosha, *Drosophila* Dicer-2, *Giardia intestinalis* Dicer, and *Arabidopsis* DCL-1 and DCL-3, the Dcr-1 RNase III domains form an intramolecular dimer (Figures 2A and S6) (Zhang et al., 2004; Macrae et al., 2006; Kwon et al., 2016; Liu et al., 2018; Sinha et al., 2018; Wang et al., 2021; Wei et al., 2021). In structures IIa and IIb, the two catalytic sites flank the minor groove of the pre-miRNA and interact with both arms of the pre-miRNA stem, confirming that they correspond to cleavage-competent complexes. The conserved RNase IIIa (E1745, D1749, D1905 and E1908) and RNase IIIb (E2032, D2036, D2136 and E2139) catalytic quartets each coordinate Ca^2+^ ions in structure IIa (Figures 4A and 4B). Ca^2+^ likely neutralizes electrostatic repulsion between the enzyme and pre-miRNA, promoting stable binding but inhibiting cleavage.

**Figure 4.**
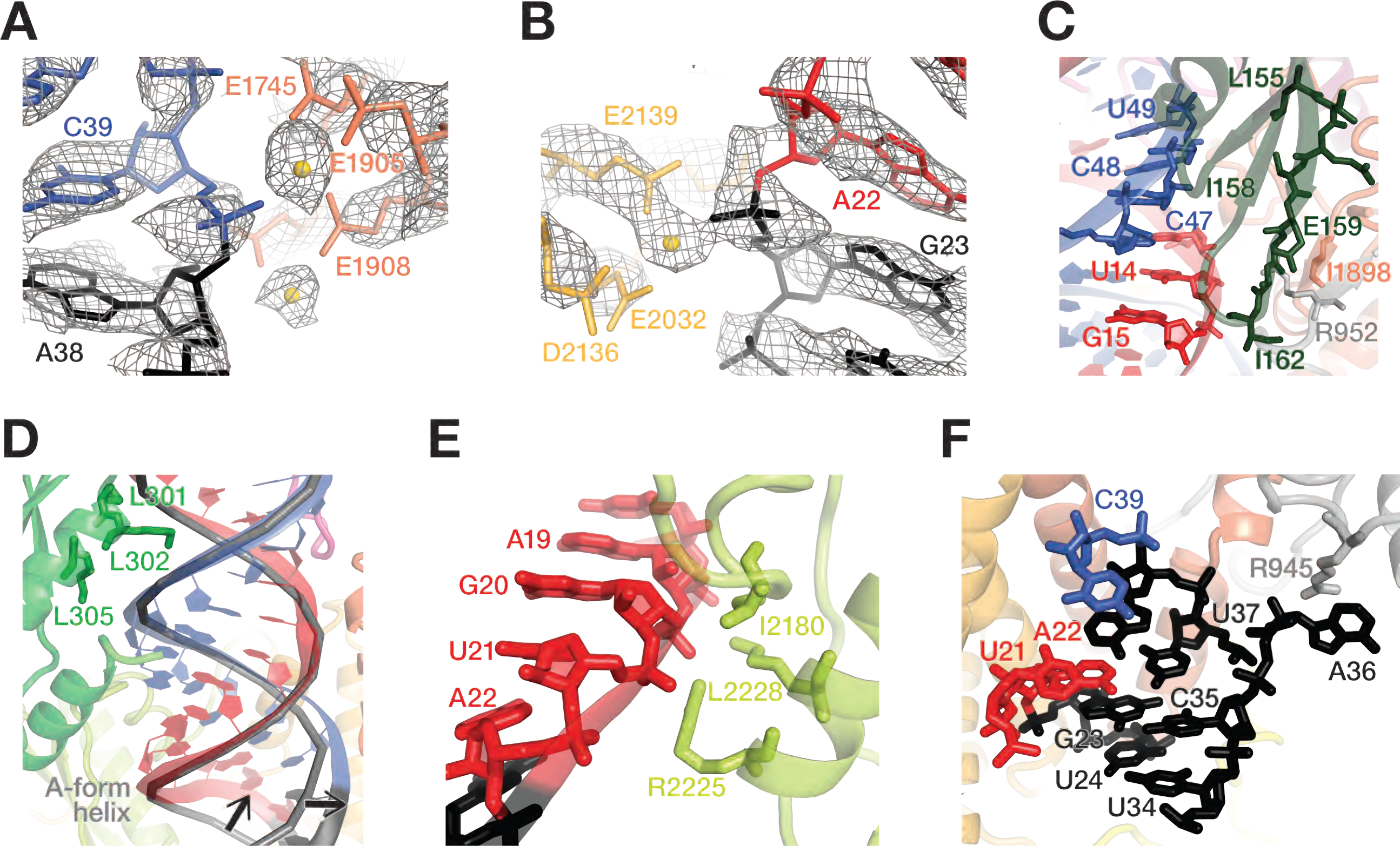
Dcr-1 and Loqs-PB position the pre-miRNA in the dicing center. **(A** and **B)** Cryo-EM density (mesh) of RNase IIIa (dark orange) and **(A)** RNase IIIb (light orange) **(B)** active site residues in the pre-miRNA-bound Dcr-1•Loqs-PB complex (structure IIa). Gold spheres: Ca^2+^ ions. **(C)** Cartoon representation of dsRBD-1 of Loqs-PB (dark green) and the Dcr-1 wing (gray and orange) stabilizing the upper part of the RNA stem. Interacting residues are shown as sticks. **(D** and **E)** Cartoon representation of dsRBD-2 of Loqs-PB **(D)** and the dsRBD of Dcr-1 **(E)** stabilizing the lower part of the pre-miRNA stem. DsRBD-2 of Loqs-PB and the Dcr-1 dsRBD are shown in green and lime green, respectively. A perfect A-form helix (predicted by RNAComposer) has been superposed as a gray ribbon. Black arrows indicate the displacement of the pre-miRNA 5′ arm by 4 Å and of the 3′ arm by 3 Å relative to their positions in an A-form helix. **(F)** Cartoon representation of the extended pre-miRNA lower stem. Adenosine A36 is bulged out and supported by a π-cation interaction with Arg945 of Dcr-1 (gray). All panels correspond to structure IIa. See also Figure S6.

Our data reveal that Loqs-PB and Dcr-1 collaborate to position the pre-miRNA into the negatively charged RNA-processing center. In structures IIa and IIb, two sets of Loqs-PB and Dcr-1 interactions stabilize the pre-cleavage state of pre-miRNA. First, dsRBD-1 of Loqs-PB is docked onto the Dcr-1 RNase IIIa (Figure 4C). Specifically, the tip (Leu155–Ile162) of the β-sheet of Loqs-PB dsRBD-1 packs against the small β-sheet formed by the Dcr-1 RNase IIIa (in the vicinity of Ile1898) and wing (in the vicinity of Arg952 and Tyr954). The interaction of the RNA-bound domain of Loqs-PB with the RNase domain of Dcr-1 stabilizes the upper part of the RNA stem near the active sites (Figure 4B). Second, the Loqs-PB dsRBD-2 and the Dcr-1 dsRBD stabilize the lower part of the pre-miRNA stem by interacting with two consecutive minor grooves of the 21- bp stem (Figure 4D). Lys301, Lys302, and Lys305 of the evolutionarily conserved KKxAK motif of the Loqs-PB dsRBD-2 recognize the phosphate backbone of the pre-miRNA stem. The 5′ arm of the stem is further positioned by positively charged Dcr-1 residues that form salt bridges with the backbone phosphates of pre-miRNA A22 (with Arg2225) and G20 (with Lys2228) and by the side chain of Dcr-1 residue Ile2180, which packs against the ribose sugar of nucleotide A19 (Figure 4E). Loop adenosine A36, which lies two nucleotides 3′ to the scissile phosphate of the 5′ arm of the stem, is bulged out and supported by a π-cation interaction with Arg945 of Dcr-1. Loop nucleotides guanosine G23 and cytosine C35 form a canonical Watson-Crick base-pair, and uracils U24 and U34 form a non-canonical Watson-Crick-like base-pair, extending the pre-miRNA stem and directing the pre-miRNA loop into the loop-binding pocket of Dcr-1 (Figure 4F). Together, these contacts position the pre-miRNA stem in the dicing center and stabilize displacement of the scissile phosphate at the 3′ end of the prospective miRNA by 4 Å and at the 5′ end of the future miRNA* by 3 Å relative to their positions in an A-form helix (shown by arrows in Figure 4D). This displacement moves the scissile phosphates into the RNase IIIb and IIIa active sites, achieving a catalytically competent geometry (Figures 4D and 5A). The conformational strain resulting from the distortion of the RNA helix is consistent with conformational proofreading accounting for cleavage specificity. Furthermore, the release of this strain after the cleavage reaction likely drives the RNA rearrangements and product release observed in structures III and IV (described below).

**Figure 5.**
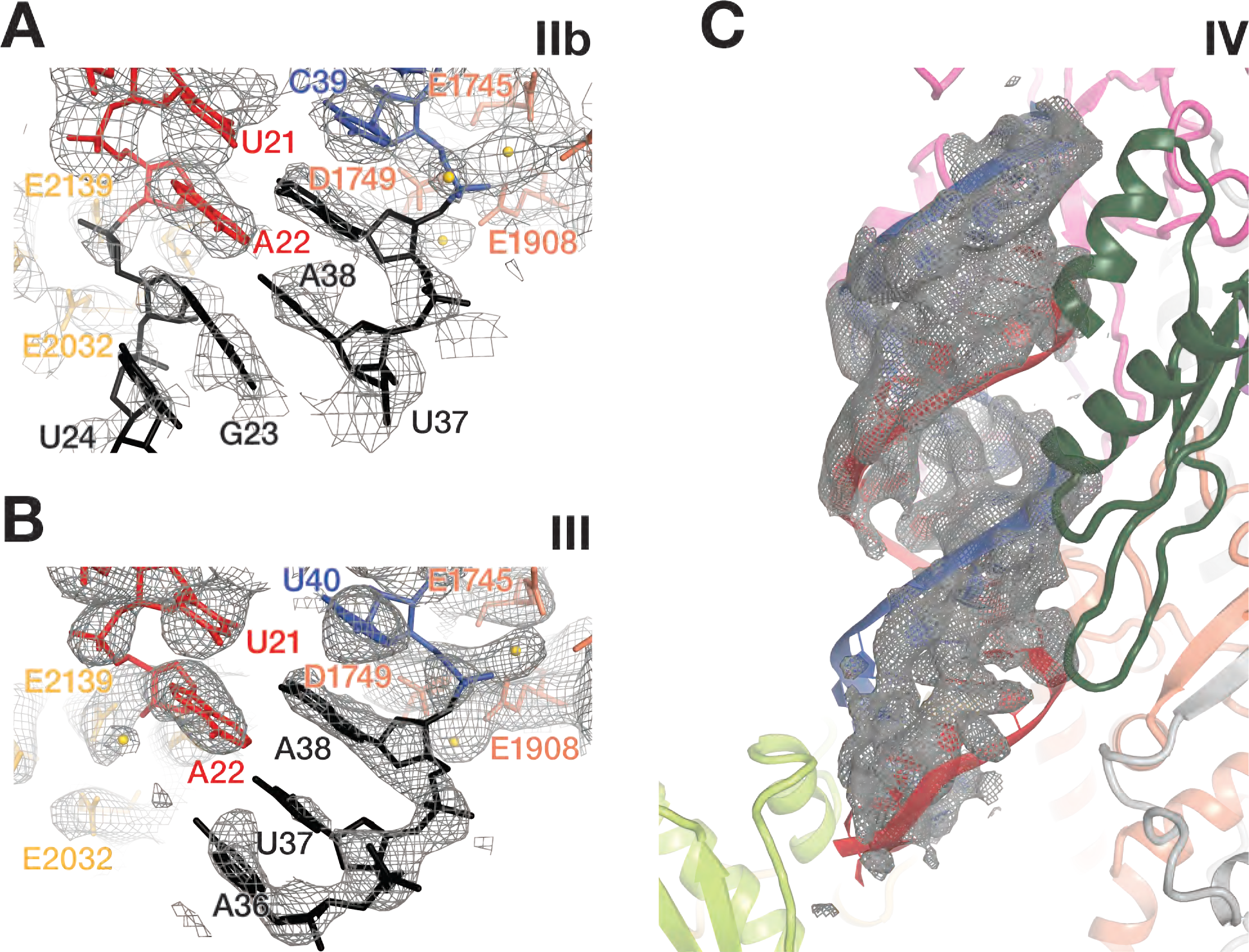
Sequential cleavage of the pre-miRNA stem. **(A** and **B)** Cryo-EM density (mesh) of RNase IIIa and RNase IIIb active centers of RNA- bound Dcr-1•Loqs-PB complex in structure IIa **(A)** and III **(B)**. The residues composing the catalytic RNase IIIa and RNase IIIb sites are shown in dark and light orange, respectively. Gold spheres: Ca^2+^ ions. **(C)** Cryo-EM density (mesh) corresponding to the miRNA/miRNA* duplex seen in structure IV, in which both arms of the pre-miRNA stem have been cleaved. Superposed is the model of Dcr-1•Loqs-PB•pre-miRNA shown in cartoon representation.

### Sequential Cleavage of the 5′ and 3′ Arms of the Pre-miRNA Stem

RNase III enzymes cleave each RNA strand via a bimolecular nucleophilic substitution mechanism that requires two Mg^2+^ ions and leaves 3′-hydroxyl and 5′-monophosphate product termini (JJ, 1982; Hutvágner et al., 2001; Campbell et al., 2002; Sun et al., 2005; Gan et al., 2008; Court et al., 2013; Nicholson, 2014). Structures IIa and IIb are consistent with this mechanism, and both the Ca^2+^ (structure IIa) and Mg^2+^ (structure IIb) structures reveal a pair of divalent cations in the RNase IIIa active site (Figures 4A and 5A). A single Ca^2+^ ion was well-resolved in RNase IIIb active site, whereas weaker density suggests that the second ion may be bound with partial occupancy (Figure 4B).

In structure III, the RNase IIIa domain binds the 3′ arm of the pre-miRNA; two Mg^2+^ ions are coordinated in the active site. Strong density indicates that the 3′ arm is intact and extends past the cleavage site between nucleotides A38 and C39. By contrast, no RNA density is observed in the RNase IIIb active site for the 5′ arm beyond nucleotide A22, suggesting that the 5′ arm has been cleaved and that the loop, still attached to the 3′ arm of the stem has left the RNase IIIb active site and is mobile (Figure 5B). Structure III therefore represents the Dcr-1•Loqs-PB heterodimer bound to a pre-miRNA in which the 3′ end of the miRNA on the 5′ arm has been defined but the 3′ arm remains to be cut. In contrast to structure III, both arms of the stem are diced in structure IV, producing a miRNA/miRNA* duplex product with a two-nucleotide, 3′- overhanging at each end (Figure 5C). Maximum-likelihood classification of the cryo-EM data set detected no structures with an intact 5′ arm and a cleaved 3′ arm, suggesting that processing of this model pre-miRNA is predominantly sequential, with the 5′ arm being cleaved first.

### Pre-miRNA Cleavage Allows Conformational Transitions that Define the Order of Product Release

Cleavage of the 5′ arm of the pre-miRNA does not alter the structure of Dcr-1 from its RNA-bound, pre-cleavage state. Similarly, the Loqs-PB dsRBD-1 and dsRBD-3 domains do not move relative to Dcr-1 between the two states, structures IIb and III (Figure 6A). By contrast, the Loqs-PB dsRBD-2, which is well resolved in structure IIa (Figure 2A) and partially resolved in IIb, is not visible in structure III (Figures 1B and 6A), suggesting that it becomes mobile upon cleavage of the 5′ arm and loses its interaction with the Dcr-1 dsRBD. The most notable rearrangement of the Dcr-1•Loqs-PB complex occurs upon cleavage of the 3′ arm (structure IV; Figure 6A). When both pre-miRNA arms have been cleaved, Dcr-1 undergoes partial closure (Figure 6B), accompanied by the departure of the Dcr-1 dsRBD from the cleft between the core and base. Closure narrows the loop-binding pocket, suggesting that this movement facilitates pre-miRNA loop dissociation. Structure IV features only the miRNA/miRNA* duplex, suggesting that the loop dissociates before the duplex (Figure 6A). The mature duplex remains anchored to the Dcr-1 PAZ and platform domains (Figure 6C), while the lower part of the duplex, with its newly formed, 5′ monophosphate and two-nucleotide, 3′ overhang, is shifted ∼5 Å away from its position in the catalytically competent state found in structures IIa and IIb (Figure 6D). The duplex adopts the A-form helix conformation, indicating that cleavage of the strained pre-miRNA likely drives the duplex from the active sites. The duplex-bound dsRBD-1 of Loqs-PB loses contact with the RNase IIIa domain, but dsRBD-1 remains bound to the PAZ and wing domains (Figure 6C). This suggests that the miRNA/miRNA* duplex may be presented to AGO for loading as part of the Loqs-PB•Dcr-1 complex; this idea remains to be tested.

**Figure 6.**
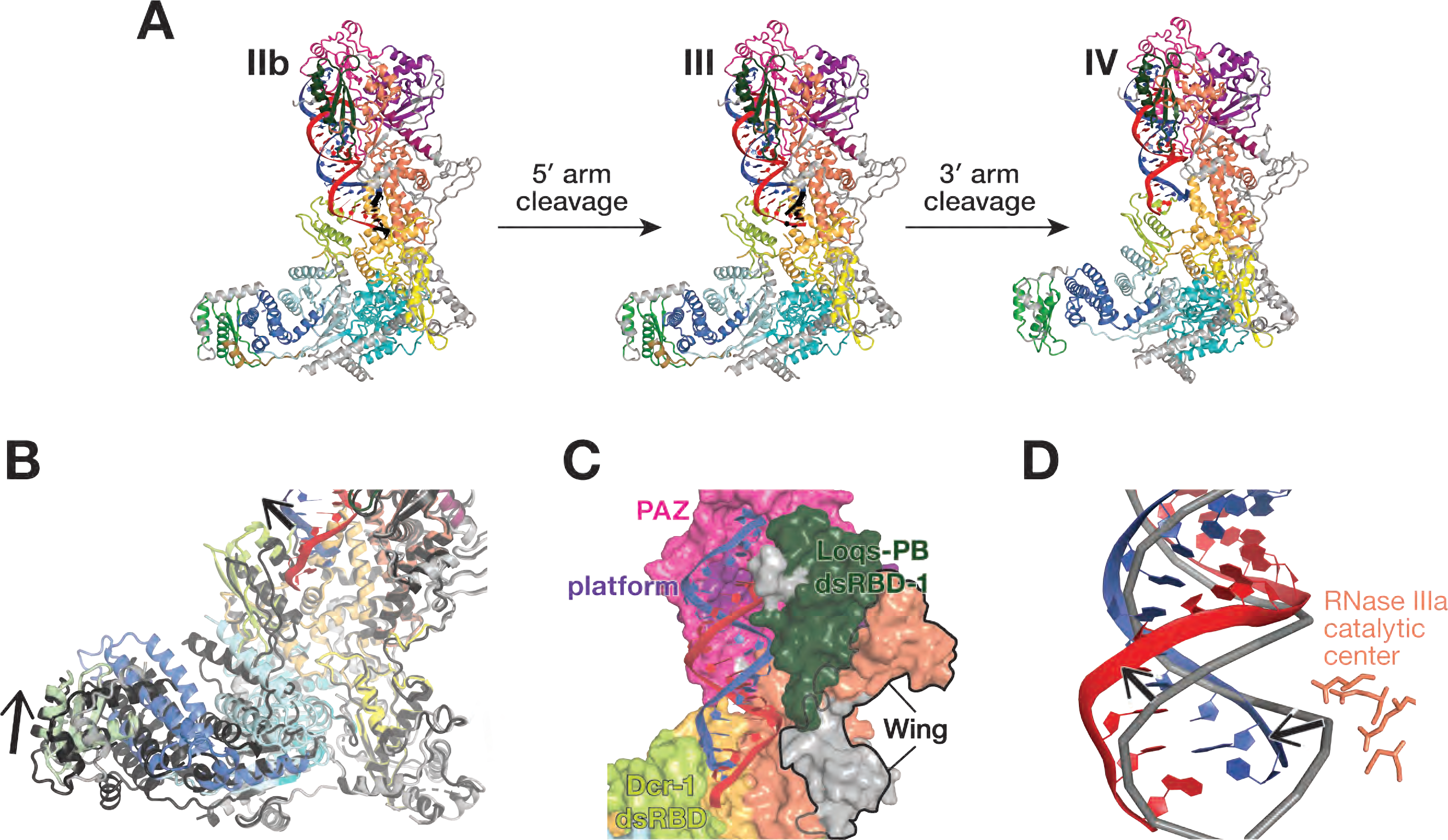
Conformational transitions define ordered product release. **(A)** Overall views of the pre-miRNA-bound Dcr-1•Loqs-PB complex in presence of Mg^2+^ before cleavage (structure IIb), after cleavage of the 5′ arm of the pre-miRNA (structure III) and after cleavage of both the 5′ and 3′ arms (structure IV). **(B)** Close-up view showing differences in the positions of the Dcr-1 dsRBD and Hel2i domains and the Loqs-PB dsRBD-3 in the Dcr-1•Loqs-PB•pre-miRNA structure IV after both 5′ and 3′ arms of the pre-miRNA have been cleaved (red and blue) relative to those in structure III after cleavage of the 5′ arm of the pre-miRNA (protein domains in dark gray). Structural alignments were performed by superposition of Dcr-1•Loqs-PB•pre- miRNA complexes. Black arrows indicate conformational changes upon the second cleavage event. **(C)** Structure of Dcr-1•Loqs-PB•pre-miRNA complex in surface (protein) and cartoon (RNA) views, showing the mature miRNA/miRNA* duplex remaining anchored to the Dcr-1 PAZ and platform domains. **(D)** Superposition of atomic models of mature duplex (structure IV) and pre-miRNA (structure IIb), showing the lower part of the duplex (red and blue) shifted ∼5 Å from its position before cleavage (gray). RNA shown in cartoon representation. RNase IIIa catalytic center (structure IV) shown as sticks.

Thus, structure IV suggests that release of the cleaved loop is a prerequisite for the subsequent departure of the miRNA/miRNA* duplex. The similarity between the Dcr-1 conformation in structure IV and the closed conformation of Dcr-1•Loqs-PB without RNA indicates that cleavage of both pre-miRNA arms and release of the loop shifts the equilibrium towards the original closed Dcr-1•Loqs-PB state (Figures 6A and 6B). This observation provides additional support for the idea that the Dcr-1•Loqs-PB heterodimer employs conformational proofreading to restrict miRNA production to authentic pre-miRNAs.

## DISCUSSION

Dicers partner with dsRBD-containing proteins in both animal and plant small RNA silencing pathways, but the mechanisms by which partner proteins enhance Dicer substrate preferences and catalytic rate is largely unknown. Our six cryo-EM structures integrate previous structural, biochemical, and biophysical observations and suggest a parsimonious model for how Dicer and its protein partners collaborate to select and process pre-miRNAs (Figure 7).

**Figure 7.**
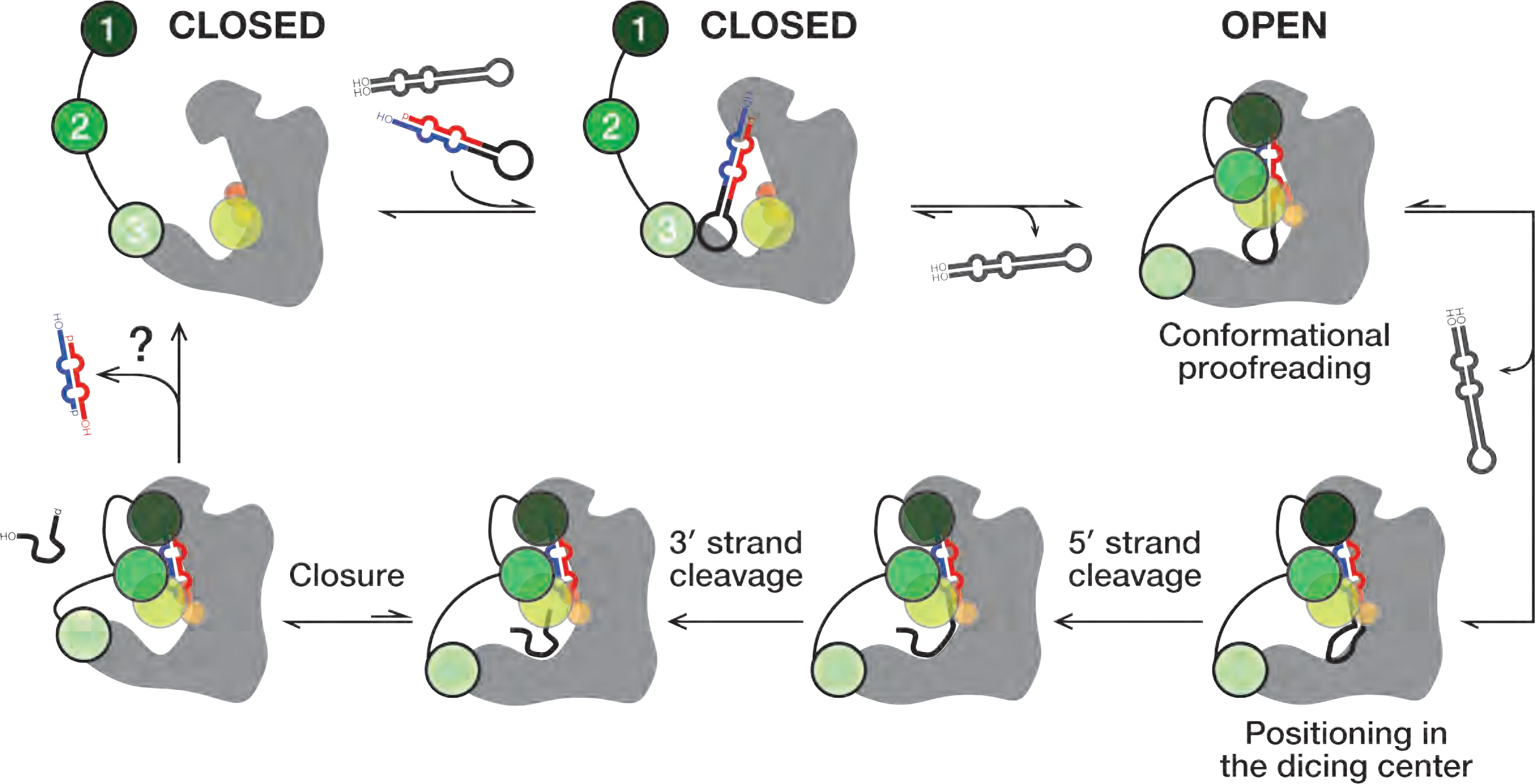
A Model for pre-miRNA processing by the Dcr-1•Loqs-PB heterodimer. Without pre-miRNA, the closed conformation of the Dcr-1•Loqs-PB complex predominates. Specific interactions between an authentic pre-miRNA and the heterodimer stabilize an open, catalytically competent state. RNA hairpins lacking the pre-miRNA structural features bind weakly and dissociate without cleavage, allowing the heterodimer to return to its closed state. The Dcr-1•Loqs-PB complex envelopes the pre-miRNA. Contacts between RNA and protein residues stabilize a distorted form of pre-miRNA, docking the RNA substrate within the Dcr-1 catalytic centers. Dcr-1 cleaves pre-miRNA sequentially. For the model-pre-miRNA used here, cleavage of the pre- miRNA 5′ arm occurs first. Subsequent cleavage of the 3′ arm generates a mature miRNA/miRNA* duplex, allowing the residual loop to depart: partial closure of Dcr-1 is coupled with release of the loop. The miRNA/miRNA* duplex remains anchored to the PAZ and platform domains of Dcr-1 and is released last.

Our cryo-EM analyses suggest that without pre-miRNA, the closed conformation of the Dicer-partner protein heterodimer predominates. In contrast, the interactions between an authentic pre-miRNA and the heterodimer stabilize an open, catalytically competent state. The conformational proofreading model predicts that dsRNA or RNA hairpins lacking the unique structural features that define a pre-miRNA would bind the Dicer-partner protein heterodimer too weakly to shift the equilibrium toward the open conformation. As a result, such binding would result in dissociation of non-pre-miRNAs without cleavage (Figure 7, far right). Supporting this idea, RNA hairpins lacking an authentic two-nucleotide 3′ overhanging end quickly dissociate from the mammalian Dicer•TRBP heterodimer (Fareh et al., 2016), and no dicing by Dcr-1 was observed for such substrates (Tsutsumi et al., 2011). By contrast, fluorescence data suggest that some substrate pre-miRNA molecules pass an “entry checkpoint,” resulting in longer residency times and cleavage on Dicer•TRBP (Fareh et al., 2016). Stabilization of the open Dcr-1 conformation by pre-miRNA likely represents the hypothesized checkpoint (Figure 7, top right).

Since not all bound pre-miRNA undergo cleavage (Fareh et al., 2016), an additional checkpoint was proposed that converts pre-miRNA into a catalytically competent state. Our structures demonstrate that catalytically competent pre-miRNA adopts a strained conformation deviating from the A-form helix. Pre-miRNA rearrangement into this conformation like represents the final checkpoint (Figure 7, bottom right), which together with Dcr-1 rearrangement accounts for the stringent conformational proofreading of potential RNA substrates.

Our structures, together with previous studies, suggest a step-wise mechanism of pre-miRNA processing by the Dcr-1•Loqs-PB heterodimer that relies on the recognition of all pre-miRNA-defining elements, from the termini to the loop. Prior to pre-miRNA binding, the closed Dcr-1 conformation features a narrow RNA-binding pocket between the core and base superdomains, and the Dcr-1 C-terminal dsRBD is bound near this pocket (structures Ia and Ib; Figure 7). In this conformation, pre-miRNA binding to the catalytic center is blocked, so it is likely that pre-accommodation states of the complex exist in which pre-miRNA is bound to Dcr-1•Loqs-PB in catalytically incompetent state(s). In one scenario, pre-miRNA may first interact with the RNA-binding Loqs-PB, which is tethered to Dcr-1 via dsRBD-3 in structures Ia and Ib (Figure 7, top center). dsRBD-1 and dsRBD-2 of Loqs-PB are not resolved, indicating that they are mobile and available to participate in the initial binding of RNA. In an alternative pathway, pre-miRNA could initially interact with the Dicer upon recognition of some specific RNA features by the protein.

Cryo-EM reconstruction of the human Dicer•TRBP complex with a model pre-miRNA is consistent with the second scenario (Liu et al., 2018). The study reported a catalytically incompetent complex conformation, in which pre-miRNA termini interact with PAZ while the loop is placed ∼40 Å away from its catalytically engaged position in our structures IIa and IIb. As in our closed-state structures Ia and Ib, the dsRBD of human Dicer in this catalytically incompetent conformation blocks entrance to the RNA processing center. Because dsRBD-1 and dsRBD-2 of TRBP are not visible in the human Dicer•TRBP complex, the partner protein appears to play a key role in positioning pre-miRNA substrate for cleavage rather than for initial engagement with potential RNA substrates. Some of our 2D class averages of the Dcr-1•Loqs-PB•pre-miRNA complex in presence of Ca^2+^ contain particles that resemble the human Dicer•TRBP•pre-miRNA complex. Nevertheless, they did not yield a similar 3D reconstruction, suggesting that if our data set contains pre-accommodation states, they are rare. This suggests that if the initial engagement of Dcr-1•Loqs-PB with pre-miRNA is in a catalytically incompetent state, displacement of the Dcr-1 dsRBD and relocation of the pre-miRNA are not rate-determining for RNA processing, and Dcr-1•Loqs-PB•pre-miRNA rapidly progresses to a catalytically competent state after substrate binding.

Our pre-miRNA-bound structures IIa and IIb show how the Dcr-1•Loqs-PB complex recognizes pre-miRNA and forms a belt around the substrate to stabilize pre-miRNA in the Dcr-1 catalytic centers. The Loqs-PB dsRBD-1 and the wing domain of Dcr-1 collaborate to secure both the 5′ and 3′ pre-miRNA ends within the PAZ and platform domains of Dcr-1. The second dsRBD of Loqs-PB cooperates with the shifted dsRBD of Dcr-1 to envelope the lower part of pre-miRNA stem in a positively charged tunnel. In this conformation, the Dcr-1 dsRBD is wedged between the protein’s base and core domains, stabilizing a widened pre-miRNA loop-binding pocket in an open Dcr-1 conformation.

Previous studies suggested several possible mechanisms by which Dicer enzymes define small RNA length—the 3′ counting rule (Macrae et al., 2006; MacRae et al., 2007), the 5′ counting rule (Park et al., 2011), and the loop counting rule (Gu et al., 2012). Our structures suggest that Dcr-1 recognizes both the 5′-phosphorylated nucleotide and the two-nucleotide 3′-overhang of pre-miRNA and support the previous model that Dicer combines 5′- and 3′-end recognition as a start to measure miRNA length (Macrae et al., 2006; Park et al., 2011; Tsutsumi et al., 2011; Tian et al., 2014).

The ∼80 Å distance from the 3′-end binding site of the PAZ domain to the loop-binding pocket formed by Hel2, DUF283 and dsRBD is consistent with the length requirement for a pre-miRNA substrate. Furthermore, our catalytically competent structures feature an unexpected 1-nt bulge at position 36, i.e., two nucleotides downstream of the RNase IIIa active center. This observation supports the ‘‘loop-counting model’’ which postulates that dicing of the 3′ strand is determined by the two-nucleotide distance between the site of cleavage and upstream loop or bulge structure of the pre-miRNA (Gu et al., 2012).

Further supporting this model, recent chemical probing of intact pre-miRNA in human cells showed that 45% of pre-miRNAs have a 3′ arm scissile phosphate that lies 2 nt away from a bulge or a loop (Luo et al., 2021).

The cleavage-competent conformation of the complex requires tight interactions of the substrate with Dcr-1 and Loqs-PB. Contacts between RNA and protein residues stabilize a distorted form of pre-miRNA in which the scissile phosphates are docked properly within the RNase IIIa and IIIb sites. Crystal structures of isolated RNase III- substrate complexes do not reveal similar RNA deviations from the A-helix geometry (Gan et al., 2008; Meng and Nicholson, 2008), suggesting that pre-miRNA distortion likely results from anchoring pre-miRNA ends in the PAZ and platform domains, docking of pre-miRNA loop in the loop-binding region, and stabilization of the RNA stem by dsRBDs of Dcr-1 and Loqs-PB.

Pre-miRNAs with central mismatches in the stem may bind Dcr-1 but fail to induce the conformational changes required for cleavage, resulting in unfavorably aligned nucleotides near the RNA-processing center. DsRBD-2 of Loqs-PB supports the dsRBD of Dcr-1 and increases pre-miRNA contacts with the phosphate backbone of RNA. We postulate that this duo of dsRBDs corrects the suboptimal geometry of difficult substrates by stabilizing an A-form-like conformation at the internal loop or cleavage site mismatches, allowing accommodation of the pre-miRNA in the catalytical center of Dcr-1. Supporting this model, previous studies showed that pre-miRNAs containing mismatches at the center of the stem or at the cleavage sites (e.g., miR-7, miR-307, miR-311, and *bantam*) require Loqs-PB for efficient dicing (Förstemann et al., 2005; Saito et al., 2005; Fukunaga et al., 2012; Lim et al., 2016; Zhu et al., 2018). Moreover, because mismatches can change the contour length of dsRNA (Holbrook et al., 1991; Baeyens et al., 1995; Sheng et al., 2014), stabilization of alternative pre-miRNA stem conformations by the Loqs-PB belt likely adjusts the pre-miRNA stem near the active centers, shifting the cleavage site by one or two nucleotides and leading to shorter or longer products than those made without Loqs-PB (Fukunaga et al., 2012). These results rationalize how Loqs-PB increases the binding affinity of Dcr-1 for pre-miRNA (Jiang et al., 2005) and stabilizes pre-miRNA structure for catalysis (Lim et al., 2016).

The precise positioning of the RNA substrate is followed by classical RNase III cleavage to generate diced miRNA/miRNA* duplex and a residual stem loop. Dicer cleavage is unlikely to be concerted: mutations that inactivate one of the two RNase III active sites of human Dicer still allow the other active site to nick the stem on one arm (Gurtan et al., 2012). Our structure III shows that Dcr-1 cleaves the 5′ and 3′ arms of our model pre-miRNA sequentially. Whether this order is universal or is determined by the sequence and structure of the pre-miRNA stem remains to be determined.

After cleavage (structure IV), the miRNA/miRNA* duplex remains anchored to the PAZ and platform domains of Dcr-1 (Figure 7, bottom left). The absence of density for the loop and the narrowing of the Dcr-1 loop-binding pocket indicate that the loop product has been released from the enzyme. The release of the loop before the duplex product likely reflects the relief of conformational strain after cleavage of the stem, as well as the paucity of stabilizing interactions between the loop and the Dcr-1 loop-binding pocket, which presents a neutral surface as opposed to the positively-charged stem-binding regions of Dcr-1 and Loqs-PB. Loop release is coupled with closure of Dcr-1 in structure IV, in keeping with the predominance of the closed Dicer conformations in the absence of pre-miRNA. Following release of the miRNA/miRNA* product, we envision Dcr-1•Loqs-PB closing, allowing for another round of conformational proofreading of substrate.

## ACKNOWLEDGEMENTS

We thank members of the Zamore, Korostelev and Hall laboratories for critical comments on the manuscript; Chen Xu, Kangkang Song, Christna Ouch and Kyounghwan Lee for data collection at the cryo-EM facility at UMass Medical School; Anna B. Loveland for assistance with data processing. This work was supported in part by National Institutes of Health grants R35 GM136275 (P.D.Z.) and R35 GM127094 (A.A.K.), and by the Intramural Research Program of the National Institutes of Health, National Institute of Environmental Health Sciences (1ZIA50165) to T.M.T.H. K.J. was supported by a Charles A. King Trust Postdoctoral Fellowship.

## AUTHOR CONTRIBUTION

Conceptualization: T.M.T.H., P.D.Z. and A.A.K. Methodology and investigation: K.J., D.G., R.C.D. and G.D. Data analyses: K.J., D.G., and A.A.K. Resources: K.J. and R.C.D. Writing-original draft: K.J. Writing-review and editing: all; visualization: K.J. Supervision: T.M.T.H., P.D.Z., and A.A.K. Funding acquisition: K.J., T.M.T.H., P.D.Z., and A.A.K.

## STAR METHODS

### METHOD DETAILS

#### Recombinant Dcr-1 and Loqs-PB Expression and Purification

Sf9 cells were maintained in suspension in SFX-Insect serum-free media (HyClone Laboratories) with Penicillin (100 U/ml)-Streptomycin (0.1 mg/ml) (Sigma) at 27°C. His_6_- Dcr-1 and His_6_-Loqs-PB were co-expressed in Sf9 cells using the BAC-to-BAC Baculovirus Expression System (Invitrogen). Sf9 cells were co-infected with His_6_-Dcr-1 (multiplicity of infection (MOI) 5) and His_6_-Loqs-PB (MOI 2.5) viral stocks and incubated for 80–90 h at 27°C. Cells were pelleted at 500 × *g* and stored at –80°C until use. Cells were thawed on ice and re-suspended with ice-cold buffer containing 30 mM HEPES-NaOH, pH 7.4, 300 mM NaCl, 10% (w/v) glycerol, 20 mM Imidazole, 2 mM DTT, 1 mM AEBSF, hydrochloride, 0.3 μM Aprotinin, 40 μM Bestatin, hydrochloride, 10 μM E-64, 10 μM Leupeptin hemisulfate, and then lysed on ice with a Dounce homogenizer using 40 strokes of a tight pestle (B type). The homogenate was centrifuged at 500 × *g* to remove nuclei and cell membranes, then at 100,000 × *g* for 20 min at 4°C.

All chromatography was conducted at 4°C. The S100 supernatant was loaded onto a column packed with 5 ml Ni-Sepharose (GE Healthcare). The column was washed with 50 ml wash buffer (30 mM HEPES-NaOH, pH 7.4, 300 mM NaCl, 10% (w/v) glycerol, 2 mM DTT) containing 20 mM imidazole, then with 50 ml wash buffer containing 40 mM Imidazole. The Dcr-1•Loqs-PB complex was eluted with 30 ml wash buffer containing 500 mM imidazole. Fractions containing Dcr-1•Loqs-PB were pooled, adjusted to 30 mM HEPES-NaOH, pH 7.4, 100 mM NaCl, 10% (w/v) glycerol, 2 mM EDTA, 2 mM DTT, and loaded onto a column packed with 5 ml Q Sepharose (GE Healthcare). The column was washed with 50 ml of column buffer (30 mM HEPES-NaOH, pH 7.4, 10% (w/v) glycerol, 2 mM EDTA, 2 mM DTT) containing 100 mM NaCl, then with 30 ml column buffer containing 200 mM NaCl. Dcr-1•Loqs-PB complex was eluted with column buffer containing 450 mM NaCl. Fractions containing Dcr-1•Loqs-PB were pooled and diluted 4.5-fold with 20 mM HEPES-NaOH, pH 7.0, 10% (w/v) glycerol, 2 mM EDTA, 2 mM DTT, to reduce the NaCl concentration to 100 mM. The sample was loaded onto a column packed with 1 ml Heparin Sepharose (GE Healthcare). The column was washed with 10 ml of 20 mM HEPES-NaOH, pH 7.0, 100 mM NaCl, 10% (w/v) glycerol, 2 mM EDTA, 2 mM DTT, then with 20 mM HEPES-NaOH, pH 7.0, 100 mM NaCl, 10% (w/v) glycerol, 2 mM EDTA, 2 mM DTT, 1 mM ATP. Dcr-1•Loqs-PB was eluted with 20 mM HEPES-NaOH, pH 7.0, 400 mM NaCl, 10% (w/v) glycerol, 2 mM EDTA, 2 mM DTT. Fractions containing Dcr-1•Loqs-PB were pooled, adjusted to 20 mM HEPES, pH 7.0, 1.5 M NaCl, 10% (w/v) glycerol, 2 mM DTT, and loaded onto a column packed with 1 ml Phenyl agarose (GE Healthcare). The column was washed with 10 ml of buffer A (20 mM HEPES, pH 7.0, 10% (w/v) glycerol, 2 mM DTT) containing 1.5 M NaCl. Dcr-1•Loqs-PB was eluted with a 30 ml linear gradient from 1.5 M to 0 M NaCl in buffer A. Fractions containing Dcr-1•Loqs-PB were pooled together and dialyzed at 4°C against three changes (3 h each) of a 1,000-fold excess of 20 mM HEPES-KOH, pH 7.9, 100 mM KCH_3_CO_2_, 5% (w/v) glycerol, 2 mM DTT. Immediately before preparing grids, the complex was diluted progressively with 20 mM HEPES-KOH, pH 7.9, 100 mM KCH_3_CO_2_, 0.1% (w/v) glycerol, 2 mM TCEP, in five steps to reduce the glycerol concentration to 0.1% (w/v) and then reconcentrated with a 30 kDa cut-off Amicon Ultra centrifugal filter (Millipore) to 3 µM final concentration.

#### In Vitro Reconstitution of the Dcr-1•Loqs-PB•Pre-miRNA Complex

The model pre-miRNA corresponded to pre-let-7 with a fully complementary stem (5′- UGA GGU AGU AGG UUG UAU AGU AGU AAU UAC ACA UCA UAC UAU ACA ACC UAC UAC CUC UCU-3′; Sigma). Pre-miRNA was 5′-phosphorylated and gel-purified. Purified Dcr-1•Loqs-PB complex was diluted to 1 µM in 20 mM HEPES-KOH, pH 7.9, 100 mM KCH_3_CO_2_, 0.1% (w/v) glycerol, 2 mM TCEP and 3 µM either Ca(CH_3_CO_2_)_2_ or Mg(CH_3_CO_2_)_2_. RNA was added in two-fold molar excess over heterodimer. To preserve protein-RNA interactions and inhibit dicing, Dcr-1•Loqs-PB•pre-miRNA complex was assembled in the presence of Ca^2+^ on ice for 1 h. To enable dicing, all components were pre-warmed to 25°C and Dcr-1•Loqs-PB•pre-miRNA complex was assembled in the presence of Mg^2+^ at 25°C for 5 min.

#### Cryo-EM Specimen Preparation

Fresh Dcr-1•Loqs-PB and Dcr-1•Loqs-PB•pre-miRNA complexes were purified immediately before preparing the frozen-hydrated grids. Holey-carbon grids (C-flat Holey Carbon Grids CF-1.2/1.3 400 mesh, Electron Microscopy Sciences) were glow discharged at 20 mA with negative polarity for 60 s in a PELCO easiGlow glow discharge unit. A drop of 2.5 µl (Dcr-1•Loqs-PB complex) or 3 µl (Dcr-1•Loqs-PB•pre-miRNA complex) sample was applied to the grids. Grids were blotted at blotting force 10 for 5 s at 4°C (Dcr-1•Loqs-PB), or at blotting force 10 for 5 s at 4°C (Dcr-1•Loqs-PB•pre-miRNA in presence of Ca^2+^) or at 25°C (Dcr-1•Loqs-PB•pre-miRNA in presence of Mg^2+^). Grids were blotted at 100% humidity and plunged into liquid ethane using a Vitrobot MK4 (FEI). Grids were stored in liquid nitrogen.

#### Cryo-EM Data Collection for the Dcr-1•Loqs-PB Heterodimer

A data set of 2,849 movies was collected for the Dcr-1•Loqs-PB complex on a Talos Arctica microscope operating at 200 kV (UMass Chan Medial School Cryo-EM Center). Images were recorded using the defocus range from −0.5 to −3.0 μm on a K3 Summit direct electron detector (Gatan), using SerialEM (Mastronarde, 2005) Each exposure was acquired with continuous frame streaming at 30 frames per 1.618 s yielding a total dose of 48.06 e^-^/Å^2^. The nominal magnification was 45,000 and the calibrated super-resolution pixel size at the specimen level was 0.435 Å.

#### Cryo-EM Data Collection for Ternary Dcr-1•Loqs-PB•Pre-miRNA Complexes

Cryo-EM data sets were collected on a Titan Krios electron microscope (FEI) operating at 300 kV, equipped with a Gatan Image Filter (GIF; UMass Chan Medial School Cryo- EM Center). Images were recorded using the defocus range from −0.5 to −3.5 µm in the super-resolution mode of the K3 Summit direct electron detector (Gatan). During multi-shot data collection, data were recorded from two holes at a time, using SerialEM as described above. Magnifications for the native and Ca^2+^-inhibited reactions were 105,000 and 81,000, yielding the super-resolution pixel sizes of 0.415 Å and 0.53 Å, respectively. For the native reaction supplemented with Mg^2+^, a total of 8,128, 38-frame movies were recorded from two grids, with an electron dose rate of 1.75 e^-^/Å^2^ and exposure time of 1.89 s, totaling 66.3 e^-^/Å^2^. For the reaction with Ca^2+^, 3,056, 40-frame movies were collected with an electron dose rate of 1.64 e^-^/Å2 and exposure time of 2.8 s, a total dose of 65.5 e^-^/Å^2^.

### QUANTIFICATION AND STATISTICAL ANALYSIS

#### Data Processing for the Dcr-1•Loqs-PB Heterodimer

Preliminary data analysis was done in Relion 3 (beta version) (Zivanov et al., 2018), including motion-correction summing to pixel size of 0.87 Å and CTF estimation on 2,849 movies. Particle picking with crYOLO/JANNI (Wagner et al., 2019; Wagner and Raunser, 2020) using default pretrained neural network models for denoising and picking yielded 345,112 coordinates. Particles were extracted from micrographs as a 4× binned stack with the pixel size of 3.48 Å, and the stack was subjected to 2D classification in Relion into 300 classes. Particles from Dicer-like containing classes were used to generate an *ab initio* 3D map that was smoothed using the gaussian filter implemented in ChimeraX (Pettersen et al., 2021), and the 3D map was used for template particle picking in the next step.

All 2,849 movies were patch motion-corrected without Fourier cropping as implemented in cryoSPARC v3 (Punjani et al., 2020). After the “Patch CTF” estimation, images with excessive total motion of the sample or CTF fit lower than 10 Å were removed, and further processing was done on 2,780 micrographs. The *ab initio* map obtained using Relion was used to create 50 equally spaced 2D templates, and both templates and micrographs were low-pass-filtered to 16-Å resolution. Template particle picking with the particle diameter set to 180 Å and the minimal separation distance set to 0.5 Å, yielded 1,478,681 particles, which were extracted as 98-px boxes with pixel size 5.2 Å. This set of particles was subjected to 2D classification into 200 classes using a 200-Å circular mask. After discarding particles that belonged to low-resolution or non-particle classes, 654,052 particles were re-extracted as a new stack with the box size of 160 px and the pixel size of 1.74 Å, and the particles were subjected to 2D classification into 80 classes. After discarding lowest-resolution and featureless classes, 615,013 particles were exported for the further analysis as two separate stacks with the pixel sizes of 0.87 Å and 3.48 Å. We performed two independent 3D classifications of the 3.48 Å/px stack into 10 classes with two types of masks. First, classification with a general 3D mask on the heterodimer revealed positional flexibility of the head relative to the core and base. Second, classification using a 33-Å radius mask covering dsRBD of Dcr-1 revealed different features of dsRBD density, suggesting conformational heterogeneity of dsRBD near its binding sites in maps Ia and Ib, where dsRBD density was most interpretable. Particles corresponding to classes Ia and Ib also featured the best resolved linker between RNase IIIb and dsRBD. They were selected from the stack of pixel size 0.87 Å as 91,617- and 112,398-particle substacks, and were subjected to non-uniform refinement in cryoSPARC resulting in 4.0-Å and 3.9-Å final maps.

#### Data Processing for Dcr-1•Loqs-PB•Pre-miRNA with Mg^2+^

The data set from the first grid, containing 4,015 movies, was aligned in IMOD (Kremer et al., 1996) with the gain reference producing image sums with pixel size 0.83 Å. After inspection, 149 aligned images were discarded due to poor quality. Aligned 4×-binned images were denoised using the noise2noise algorithm implemented in JANNI (Wagner and Raunser, 2020) and used for initial particle picking with crYOLO (Wagner et al., 2019) with the pretrained general model, using box size 140 Å. After selecting 134,417 particle coordinates, all subsequent processing steps were performed in Relion (Zivanov et al., 2018). A 4×-binned particle stack with the pixel size of 3.32 Å was extracted from 3,866 motion-corrected and CTF-estimated micrographs, and template-free 2D classification into 168 classes was performed. Particles from nine, high-resolution classes totaling 57,184 particles were used to calculate an *ab initio* 3D map that was auto-refined using a mask against the same stack until 6.64 Å Nyquist-limit resolution. The resulting map revealed pre-miRNA bound to Dcr-1 with additional density that was attributed to one of the Loqs domains bound to pre-miRNA. This map was low-pass filtered to 20 Å, and with 30° angular sampling, 48 2D reprojections were generated as reference images for template-based particle picking.

For template-based particle picking, 8,128 movies from both grids were motion-corrected with a GPU-accelerated version of MotionCor2 (Zheng et al., 2017) to the final pixel size of 0.83 Å and defocus values were determined using CTFFIND4 (Rohou and Grigorieff, 2015). After discarding poor-quality micrographs, 7,895 aligned images were used for template-based particle picking, yielding a 3,161,540-particle stack. Boxes (360 px × 360 px) with particle coordinates were extracted as a 4×-binned stack with pixel size 3.32 Å, which was subjected to reference-free 2D classification into 386 classes for 30 iterations. Fourteen non-junk 2D classes accounted for 1,148,974 particles, which were selected for another round of 2D classification into 60 classes for 35 iterations to further improve the alignment parameters and to discard lower-quality particles. Sixteen 2D classes comprising 565,607 particles were selected for the masked 3D auto-refinement to calculate the updated 3D map. The refinement was performed with a starting map low-pass filtered to 30 Å, using a 10-Å hard-edge and a 10-Å soft-edge mask and converging to the final Nyquist-limit resolution of 6.64 Å.

The particle stack was 3D-classified using the updated 3D map low-pass filtered to 15 Å as a mask. Classification into six classes revealed two high-quality classes that differed in the region corresponding to pre-miRNA loop, however the remaining classes suggested unresolved conformational heterogeneity necessitating additional classification (described below). One of the two classes featured strong RNA-loop density, whereas the second class contained no density in the corresponding region.

Particles from both classes were re-extracted as separate 1× stacks comprising 185,002 and 106,904 particles, and the stacks refined in cryoSPARC to result in final maps IIa and IV at 3.3- and 4.0-Å resolutions, respectively.

To additionally resolve the heterogeneity and potentially separate the cleavage intermediates, we performed 3D classification of the 4×-binned stack into 10 classes without alignment with a 33-Å 3D mask centered at the RNase IIIb cleavage site (near pre-miRNA nucleotide 22). An additional well-resolved class was obtained that featured continuous density for the post-cleavage site fragment of the 3′ strand but not for the 5′ strand. Particles (135,303) were re-extracted into a 1× stack and refined in cryoSPARC with non-uniform refinement to result in map-III at 3.3-Å resolution.

#### Data processing for Dcr-1•Loqs-PB•Pre-miRNA with Ca^2+^

Movies were motion-corrected with the GPU accelerated version of MotionCor2 (Zheng et al., 2017) to the final pixel size of 1.06 Å, and defocus values were determined using CTFFIND4 (Rohou and Grigorieff, 2015) The best 2,999 aligned images were selected for further analysis. Forty-eight 2D reprojections from Gaussian low-pass filtered map to 20 Å from our previous reconstruction were used for template-based particle picking, yielding 2,023,692 particles. Boxes (282 px × 282 px) with particle coordinates were extracted into a 3×-binned stack with pixel size 3.18 Å and subjected to reference-free 2D classification into 386 classes for 20 iterations. The six most well-resolved 2D classes representing different particle orientations were selected, totaling 1,109,821 particles. A 3D classification into six classes with a low-pass filtered (20 Å resolution) *de novo* 3D reconstruction as a reference was used to select particles with the highest-resolution features. Particles (353,726) were re-extracted with box size 340 px × 340 px with re-centering and were resampled to pixel size 1.24 Å to speed up subsequent processing steps. After masked auto-refinement, CTF-refinement and Bayesian particle polishing were performed, each followed by a round of auto-refinement and postprocessing, resulting in a 3.3 Å reconstruction.

To further improve the quality of the reconstruction, the 353,726-particle stack with pixel size 1.24 Å was 3D classified without alignment into six classes, using a 3D mask created from the previous reconstruction that was low-pass filtered to 15 Å resolution and to which 3 px of hard and soft edges were added. The highest-resolution class comprised 149,443 particles that were re-extracted as a 1.06 Å/px stack and subjected to non-uniform refinement in cryoSPARC, yielding the final 3.0 Å map corresponding to structure IIa.

#### Model Building and Structure Refinement

A locally installed AlphaFold version (Jumper et al., 2021) was used to obtain the starting models of Dcr-1 and Loqs-PB. AlphaFold-Multimer (Evans et al., 2021) was used to co-fold the heterodimer of Dcr-1 helicase domains (aa 1–615) with the third dsRBD domain of Loqs-PB (aa 320–465). Dcr-1 and Loqs-PB models were rigid-body refined against the maps in ChimeraX (Pettersen et al., 2021) and COOT (Emsley et al., 2010). To this end, twelve Dcr-1 domain fragments and three Loqs-PB domains were fitted independently. Pre-miRNA nucleotides 1–24 and 34–60 were modeled in COOT, using map-IIa. Since the lower-resolution density for the RNA loop did not allow for unambitious nucleotide placement, an RNAfold/RNAcomposer-predicted (Lorenz et al., 2011; Antczak et al., 2016) starting model of the loop was generated. Secondary-structure restraints for the G23:C35 and U24:U34 pairs were used during model minimization in RNAcomposer. After the initial model building, structures of Dcr-1•Loqs-PB (Ia and Ib) and Dcr-1•Loqs-PB•pre-miRNA (IIa, IIb, III, and IV) were refined in Phenix (Adams et al., 2010) using secondary-structure restraints, and the quality of the resulting structures was assessed with MolProbity (Chen et al., 2010). The total buried surface areas at the protein-protein or protein-RNA interfaces were calculated with NACCESS (Hubbard et al., 1993). Figures were prepared with PyMol (https://pymol.org).

## SUPPLEMENTAL FIGURES

### Supplemental Figure Legends

**Figure S1.**
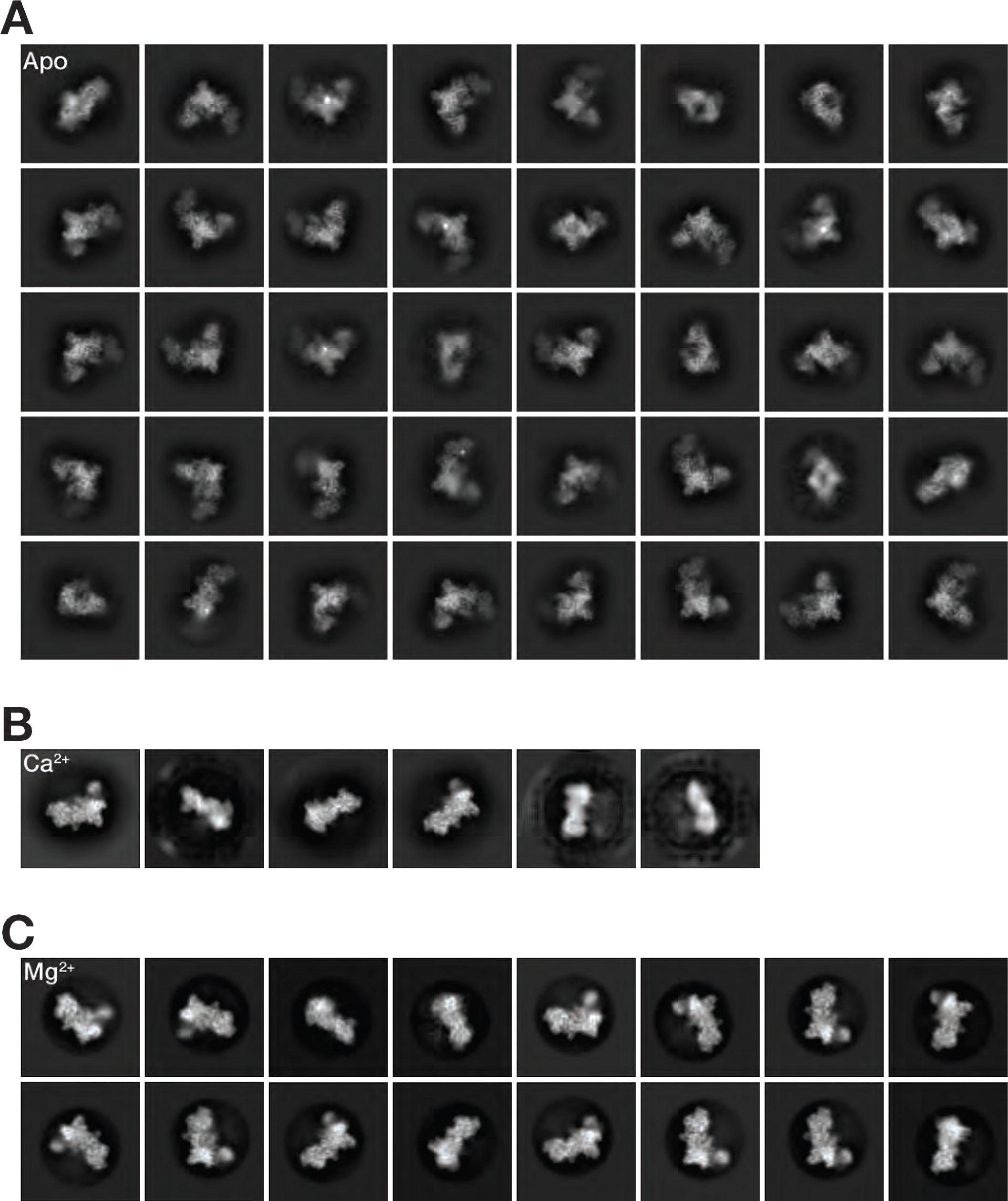
2D cryo-EM classes for the three Dcr-1•Loqs-PB data sets, related to Figure 1. **(A)** Representative 2D classes for the Dcr-1•Loqs-PB complex formed without pre-miRNA. **(B)** Representative 2D classes for the Dcr-1•Loqs-PB•pre-miRNA complex formed with Ca^2+^. **(C)** Representative 2D classes for the Dcr-1•Loqs-PB•pre-miRNA complex formed with Mg^2+^.

**Figure S2.**
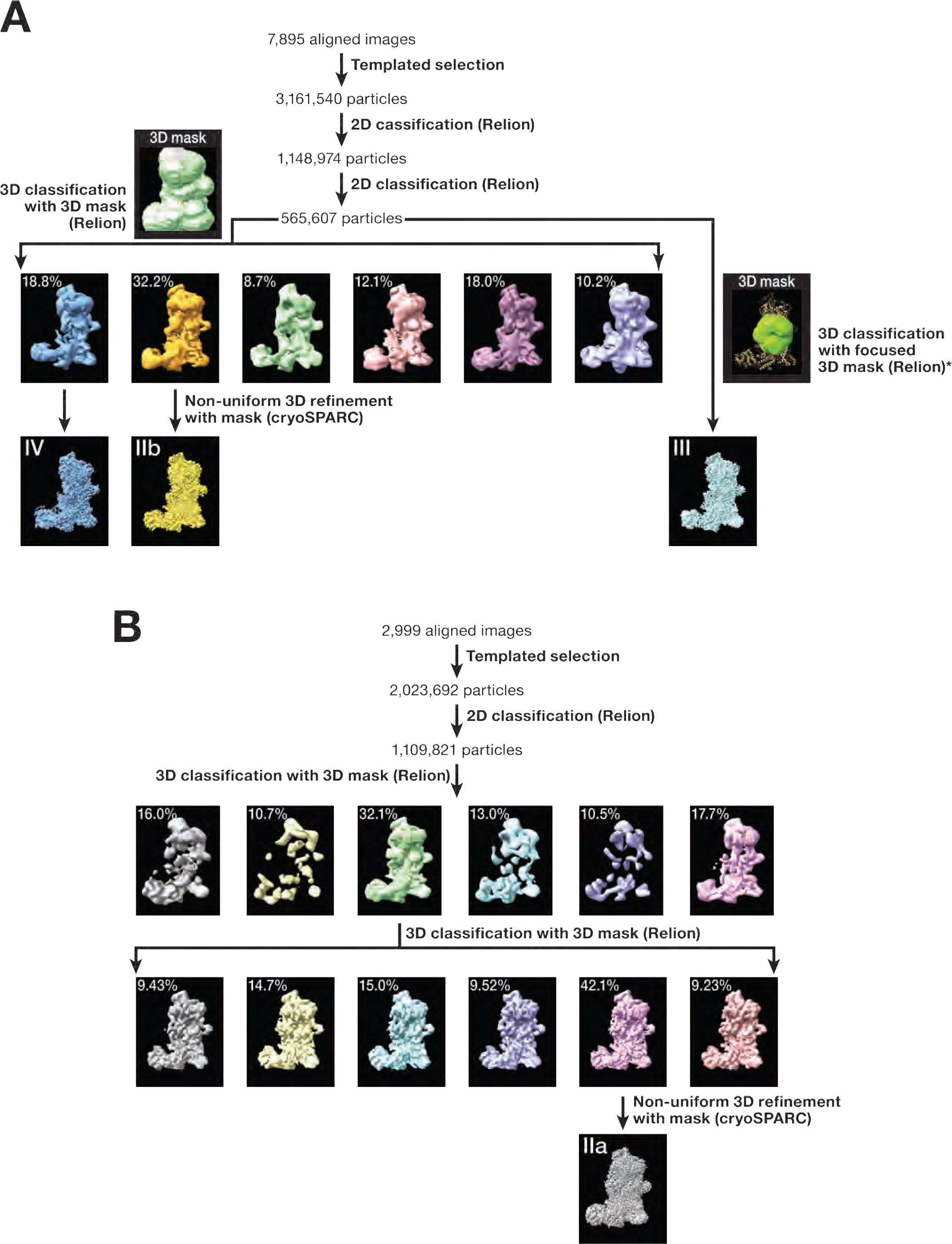
Schematics of maximum likelihood classification, related to Figure 1. **(A)** Classification of the data set for the Dcr-1•Loqs-PB•pre-miRNA complex formed with Mg^2+^. Structures IIa and IV resulted from the classification with the large mask shown on the left; structure III resulted from a similar classification using the mask centered at the Dcr-1 catalytic center (shown with *), as described in Methods. **(B)** Classification of the data set for the Dcr-1•Loqs-PB•pre-miRNA complex formed with Ca^2+^. The apo data set was processed similarly to **(A)**, with differences detailed in Methods.

**Figure S3.**
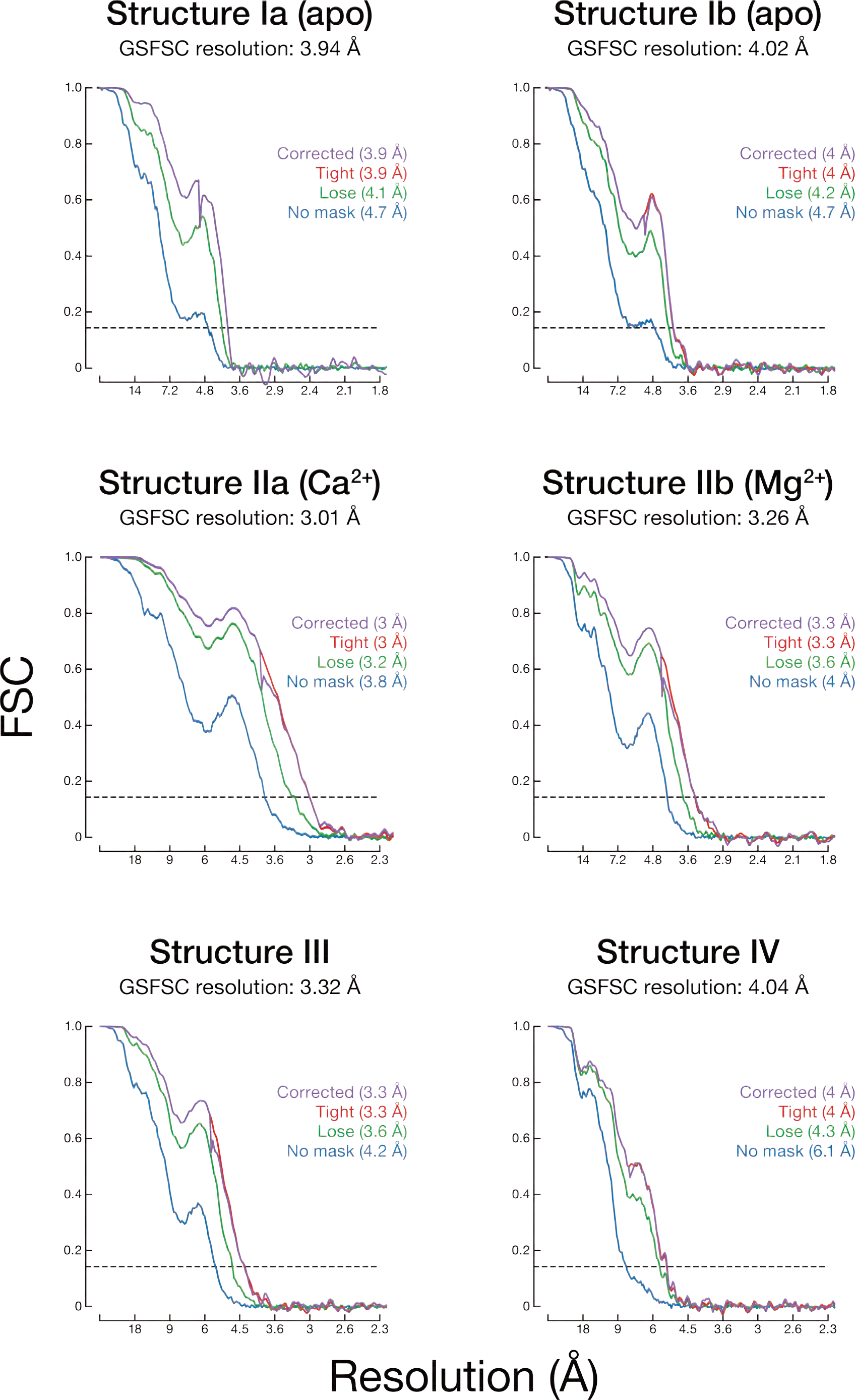
Fourier shell correlation (FSC) curves for the six cryo-EM maps resulting from classification of the three data sets by cryoSPARC, related to Figure 1. FSC curves are shown for: structures Ia and Ib obtained from the Dcr-1•Loqs-PB complex formed without pre-miRNA; Structure IIa of the Dcr-1•Loqs-PB•pre-miRNA complex formed with Ca^2+^; Structures IIb, III and IV, obtained from the data set of the Dcr-1•Loqs-PB•pre-miRNA complex formed with Mg^2+^.

**Figure S4.**
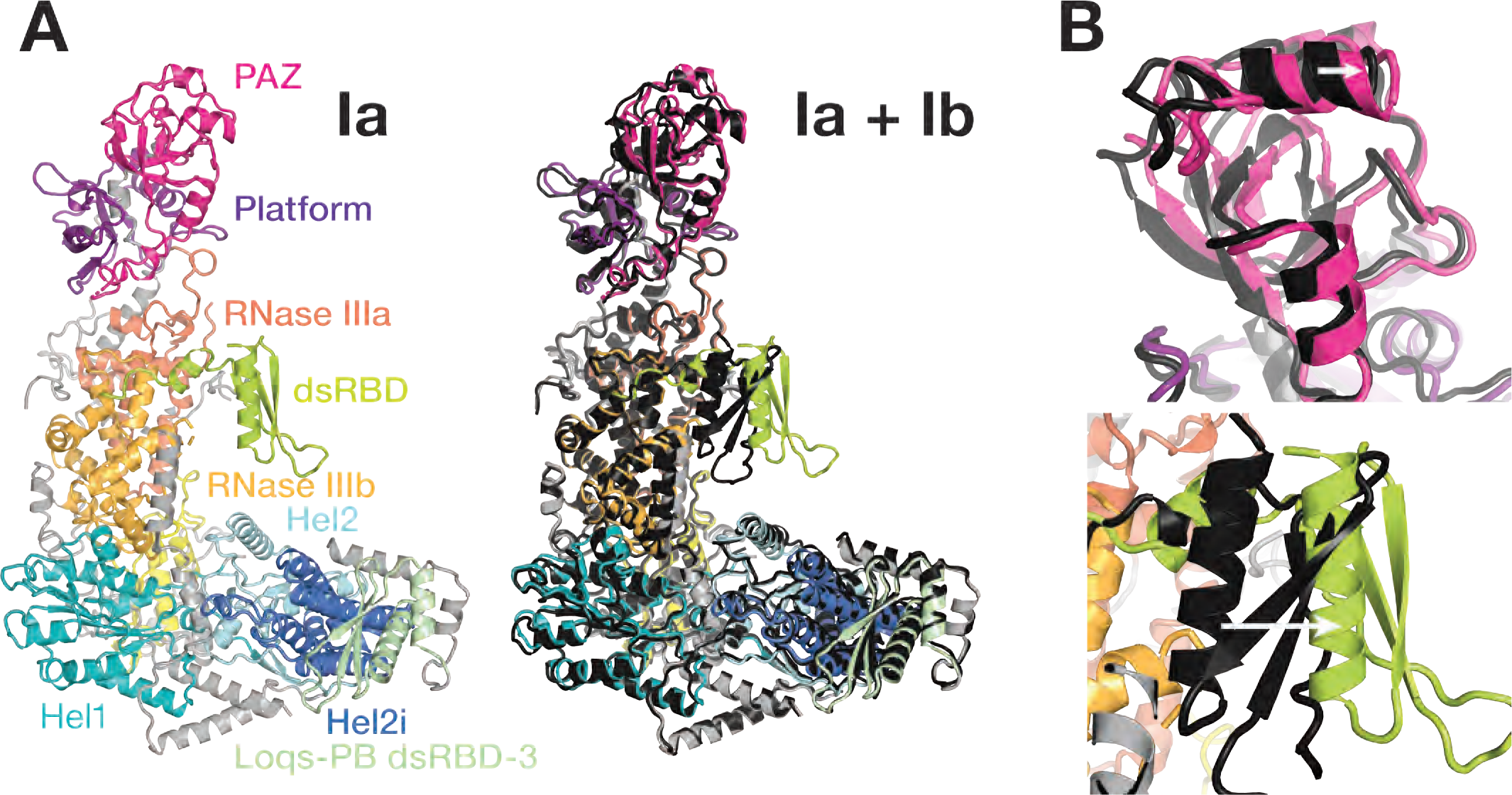
Structures of RNA-free Dcr-1•Loqs-PB heterodimer, related to Figure 2. **(A)** Overall view of the RNA-free Dcr-1•Loqs-PB complex in structures Ia and Ib, which differ by ∼10 Å and ∼3 Å movements of the Dcr-1 dsRBD and head superdomain, respectively. **(B)** Close-up views showing differences between structures Ia and Ib in the positions of the Dcr-1 dsRBD and head superdomain. Structure Ia is in color; structure Ib is shown in dark gray.

**Figure S5.**
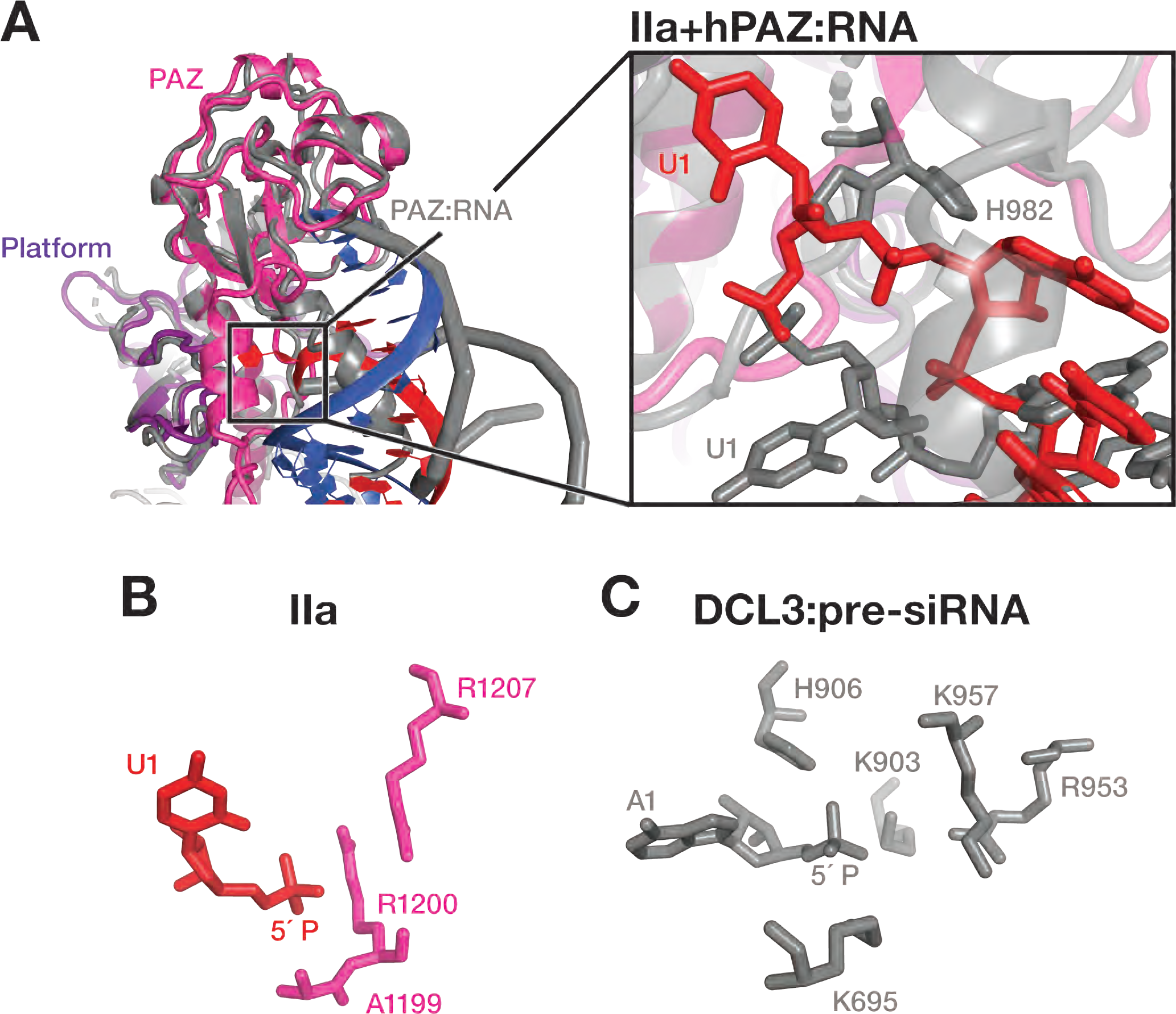
Recognition of 5′ End of pre-miRNA by Dicers, related to Figure 3. **(A)** Superposition of atomic models of RNA-bound Dcr-1•Loqs-PB complex (structure IIa) and human PAZ•RNA (structure 4NH3; {Tian et al., 2014, #94263}) in cartoon representation. Nucleotides and His982 are also shown as sticks. RNA-bound Dcr-1•Loqs-PB complex is in color; human Dicer PAZ in complex with RNA is shown in gray. **(B and C)** Comparison of the mechanism of 5′ phosphate recognition used by *D. melanogaster* Dcr-1 **(B)** and *Arabidopsis thaliana* DCL3 **(C)**. The first nucleotide of the pre-miRNA (U1) or pre-siRNA (A1) is shown with the Dicer residues involved in 5′ phosphate recognition.

**Figure S6.**
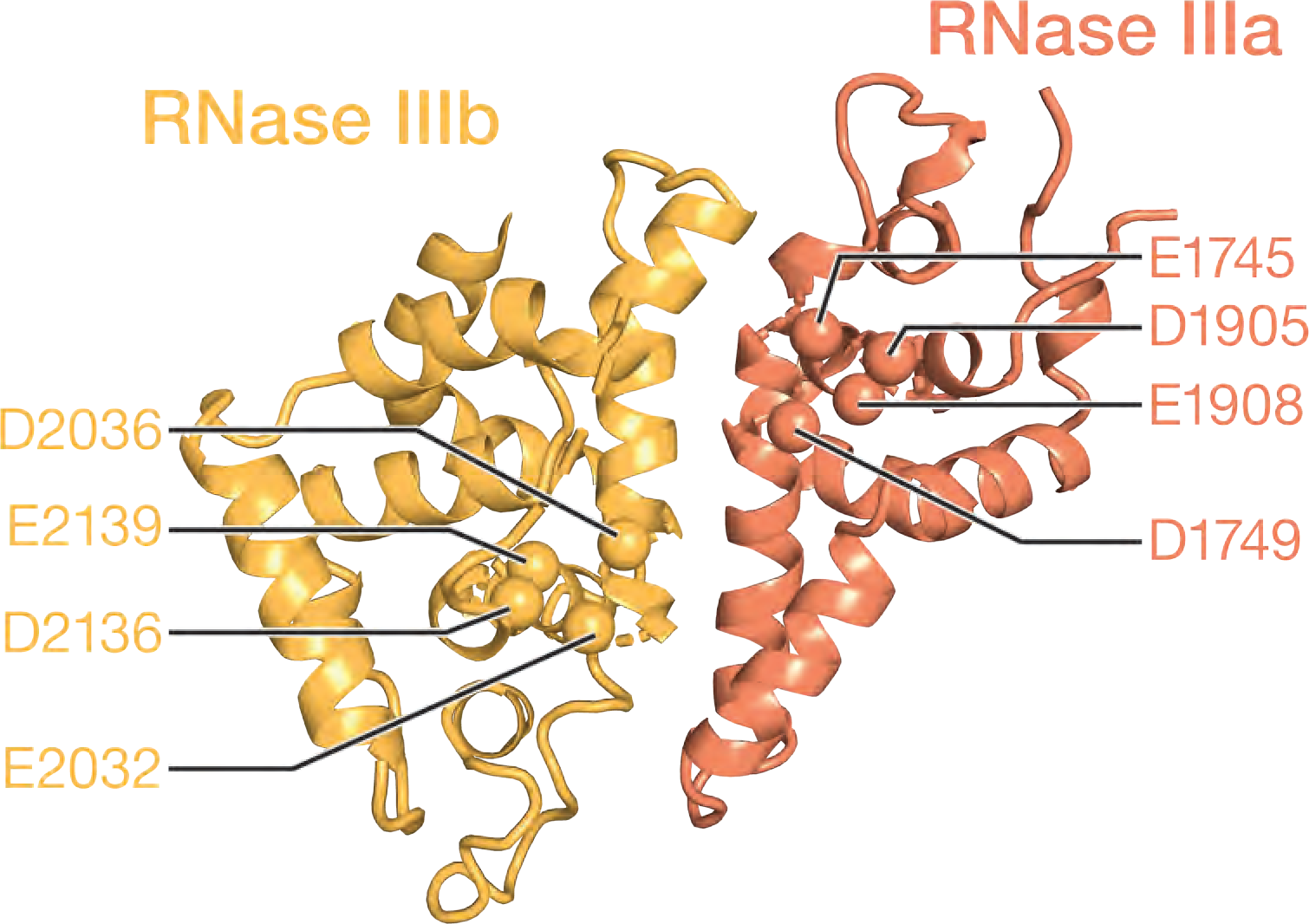
Dcr-1 RNase III domains form an intramolecular dimer, related to Figure 4. Cartoon model of Dcr-1 RNase III intramolecular dimer. Cα atoms of residues composing the catalytic sites are labeled as spheres.

**Table S1.** Cryo-EM data collection, refinement, and validation statistics.

|  | Dcr-1•Loqs-PB |  | Dcr-1•Loqs-PB•pre-miRNA |  |  |  |
| --- | --- | --- | --- | --- | --- | --- |
|  | Ia | Ib | IIa (Ca<sup>2+</sup>) | IIb (Mg<sup>2+</sup>) | III (Mg<sup>2+</sup>) | IV (Mg<sup>2+</sup>) |
| **Data collection and processing** |  |  |  |  |  |  |
| Magnification | 57,471× | 57,471× | 81,000× | 105,000× | 105,000× | 105,000× |
| Voltage (kV) | 200 | 200 | 300 | 300 | 300 | 300 |
| Electron exposure (e<sup>-</sup>/Å<sup>2</sup>) | 48.06 | 48.06 | 65.5 | 66.3 | 66.3 | 66.3 |
| Defocus range (μm) | 0.5–3 | 0.5–3 | 0.5–3.5 | 0.5–3.5 | 0.5–3.5 | 0.5–3.5 |
| Pixel size (Å) | 0.87 | 0.87 | 1.06 | 0.83 | 0.83 | 0.83 |
| Symmetry imposed | C1 | C1 | C1 | C1 | C1 | C1 |
| Initial particle images | 615,013 | 615,013 | 1,109,821 | 565,607 | 565,607 | 565,607 |
| Final particle images | 112,398 | 91,617 | 149,443 | 185,002 | 135,303 | 106,904 |
| Map resolution (Å)** | 4.02 | 3.94 | 3.01 | 3.26 | 3.32 | 4.04 |
| FSC threshold | 0.143 | 0.143 | 0.143 | 0.143 | 0.143 | 0.143 |
| **Refinement** |  |  |  |  |  |  |
| Partial model used | AlphaFold | AlphaFold | AlphaFold/<br>RNAfold | AlphaFold/<br>RNAfold | AlphaFold/<br>RNAfold | AlphaFold/<br>RNAfold |
| Model resolution (Å)* | 4.04 (7.66) | 4.06 (7.30) | 3.03 (3.52) | 3.32 (3.60) | 3.31 (3.86) | 3.71 (7.49) |
| FSC threshold | 0.143 (0.5) | 0.143 (0.5) | 0.143 (0.5) | 0.143 (0.5) | 0.143 (0.5) | 0.143 (0.5) |
| Map-sharpening *B* factor (Å<sup>2</sup>) | −93 | −89 | −111 | −117 | −120 | −141 |
| **Model composition\*** |  |  |  |  |  |  |
| Protein residues | 1690 | 1689 | 1919 | 1815 | 1815 | 1829 |
| RNA residues |  |  | 60 | 51 | 48 | 40 |
| Ions |  |  | 4 Ca<sup>2+</sup> | 11 Mg<sup>2+</sup> | 9 Mg<sup>2+</sup> |  |
| ***B* factors (Å<sup>2</sup>)\*** |  |  |  |  |  |  |
| Protein | 150 | 125 | 52.4 | 76.62 | 99.21 | 120 |
| RNA |  |  | 27.8 | 60.82 | 66.93 | 115 |
| Ions |  |  | 28.2 | 46.43 | 40.38 |  |
| **R.M.S. deviations\*,\$** | | | | | | |
| Bond lengths (Å) | 0.005 | 0.005 | 0.004 | 0.009 | 0.014 | 0.006 |
| Bond angles (degrees) | 1.050 | 1.057 | 0.760 | 1.261 | 1.376 | 1.079 |
| **Validation\*** |  |  |  |  |  |  |
| MolProbity score | 1.81 | 1.85 | 2.46 | 1.82 | 1.86 | 1.88 |
| Clashscore | 7.38 | 8.01 | 6.52 | 6.85 | 7.40 | 7.95 |
| Poor rotamers (%) | 0 | 0 | 7.46 | 0.25 | 0.06 | 0 |
| **Ramachandran plot\*** |  |  |  |  |  |  |
| Favored (%) | 93.86 | 93.81 | 93.11 | 93.25 | 92.86 | 92.95 |
| Allowed (%) | 5.96 | 5.95 | 6.52 | 6.52 | 6.91 | 6.61 |
| Disallowed (%) | 0.18 | 0.24 | 0.37 | 0.22 | 0.22 | 0.44 |
| **Validation (RNA)\*** |  |  |  |  |  |  |
| Good sugar pucker (%) |  |  | 100 | 100 | 100 | 100 |
| Good backbone of<br>overhangs + stem (%)# |  |  | 86.7 | 88.89 | 86.7 | 77.5 |

\*from Phenix; \$root mean square deviations; #RNA backbone suites that fall into recognized rotamer conformations defined by Molprobity

## REFERENCES

Adams, P. D., Afonine, P. V., Bunkóczi, G., Chen, V. B., Davis, I. W., Echols, N., Headd, J. J., Hung, L. W., Kapral, G. J., Grosse-Kunstleve, R. W., McCoy, A. J., Moriarty, N. W., Oeffner, R., Read, R. J., Richardson, D. C., Richardson, J. S., Terwilliger, T. C., and Zwart, P. H. (2010). PHENIX: a comprehensive Python-based system for macromolecular structure solution. Acta Crystallogr D Biol Crystallogr 66, 213–221.

Ambros, V., Bartel, B., Bartel, D. P., Burge, C. B., Carrington, J. C., Chen, X., Dreyfuss, G., Eddy, S. R., Griffiths-Jones, S., Marshall, M., Matzke, M., Ruvkun, G., and Tuschl, T. S. (2003). A uniform system for microRNA annotation. RNA 9, 277–279.

Antczak, M., Popenda, M., Zok, T., Sarzynska, J., Ratajczak, T., Tomczyk, K., Adamiak, R. W., and Szachniuk, M. (2016). New functionality of RNAComposer: an application to shape the axis of miR160 precursor structure. Acta Biochim Pol 63, 737–744.

Baeyens, K. J., De Bondt, H. L., and Holbrook, S. R. (1995). Structure of an RNA double helix including uracil-uracil base pairs in an internal loop. Nat. Struct. Biol. 2, 56–62.

Bartel, D. P. (2018). Metazoan MicroRNAs. Cell 173, 20–51.

Brennecke, J., Stark, A., Russell, R. B., and Cohen, S. M. (2005). Principles of microRNA-target recognition. PLoS Biol. 3, e85.

Campbell, F. E., Cassano, A. G., Anderson, V. E., and Harris, M. E. (2002). Pre-steady-state and stopped-flow fluorescence analysis of Escherichia coli ribonuclease III: insights into mechanism and conformational changes associated with binding and catalysis. J. Mol. Biol. 317, 21–40.

Cenik, E., Fukunaga, R., Lu, G., Dutcher, R., Wang, Y., Tanaka Hall, T., and Zamore, P. (2011). Phosphate and R2D2 Restrict the Substrate Specificity of Dicer-2, an ATP-Driven Ribonuclease. Mol. Cell. 42, 172–184.

Chen, S., Liu, W., Naganuma, M., Tomari, Y., and Iwakawa, H. O. (2022). Functional specialization of monocot DCL3 and DCL5 proteins through the evolution of the PAZ domain. Nucleic Acids Res. gkac223.

Chen, V. B., Arendall, W. B., Headd, J. J., Keedy, D. A., Immormino, R. M., Kapral, G. J., Murray, L. W., Richardson, J. S., and Richardson, D. C. (2010). MolProbity: all-atom structure validation for macromolecular crystallography. Acta Crystallogr D Biol Crystallogr 66, 12–21.

Chen, X. (2009). Small RNAs and Their Roles in Plant Development. Annual Review of Cell and Developmental Biology 25, 21–44.

Chendrimada, T. P., Gregory, R. I., Kumaraswamy, E., Norman, J., Cooch, N., Nishikura, K., and Shiekhattar, R. (2005). TRBP recruits the Dicer complex to Ago2 for microRNA processing and gene silencing. Nature 436, 740–744.

Chiang, H. R., Schoenfeld, L. W., Ruby, J. G., Auyeung, V. C., Spies, N., Baek, D., Johnston, W. K., Russ, C., Luo, S., Babiarz, J. E., Blelloch, R., Schroth, G. P., Nusbaum, C., and Bartel, D. P. (2010). Mammalian microRNAs: experimental evaluation of novel and previously annotated genes. Genes Dev. 24, 992–1009.

Court, D. L., Gan, J., Liang, Y. H., Shaw, G. X., Tropea, J. E., Costantino, N., Waugh, D. S., and Ji, X. (2013). RNase III: Genetics and function; structure and mechanism. Annu. Rev. Genet. 47, 405–431.

Daniels, S. M., Melendez-Peña, C. E., Scarborough, R. J., Daher, A., Christensen, H. S., El Far, M., Purcell, D. F. J., Lainé, S., and Gatignol, A. (2009). Characterization of the TRBP domain required for Dicer interaction and function in RNA interference. BMC Molecular Biology 10, http://dx.doi.org/10.1186/1471-2199-10-38.

Denli, A. M., Tops, B. B., Plasterk, R. H., Ketting, R. F., and Hannon, G. J. (2004). Processing of primary microRNAs by the Microprocessor complex. Nature 432, 231–235.

Dong, Q., Hu, B., and Zhang, C. (2022). microRNAs and Their Roles in Plant Development. Front Plant Sci 13, 824240.

JJ, D. (1982). Ribonuclease III, in The Enzymes, ed. Boyer, PD (New York: Academic Press) 15, 485-499.

Emsley, P., Lohkamp, B., Scott, W. G., and Cowtan, K. (2010). Features and development of Coot. Acta Crystallogr D Biol Crystallogr 66, 486–501.

Evans, R., O’Neill, M., Pritzel, A., Antropova, N., Senior, A., Green, T., Žídek, A., Bates, R., Blackwell, S., Yim, J., Ronneberger, O., Bodenstein, S., Zielinski, M., Bridgland, A., Potapenko, A., Cowie, A., Tunyasuvunakool, K., Jain, R., Clancy, E., Kohli, P., Jumper, J., and Hassabis, D. (2021). Protein complex prediction with AlphaFold-Multimer. bioRxiv https://doi.org/10.1101/2021.10.04.463034.

Fareh, M., Yeom, K. H., Haagsma, A. C., Chauhan, S., Heo, I., and Joo, C. (2016). TRBP ensures efficient Dicer processing of precursor microRNA in RNA-crowded environments. Nat Commun 7, 13694.

Förstemann, K., Tomari, Y., Du, T., Vagin, V. V., Denli, A. M., Bratu, D. P., Klattenhoff, C., Theurkauf, W. E., and Zamore, P. D. (2005). Normal microRNA maturation and germ-line stem cell maintenance requires Loquacious, a double-stranded RNA-binding domain protein. PLoS Biol. 3, e236.

Förstemann, K., Horwich, M. D., Wee, L., Tomari, Y., and Zamore, P. D. (2007). Drosophila microRNAs Are Sorted into Functionally Distinct Argonaute Complexes after Production by Dicer-1. Cell 130, 287–297.

Fukunaga, R., Han, B. W., Hung, J. H., Xu, J., Weng, Z., and Zamore, P. D. (2012). Dicer partner proteins tune the length of mature miRNAs in flies and mammals. Cell 151, 533–546.

Gan, J., Shaw, G., Tropea, J. E., Waugh, D. S., Court, D. L., and Ji, X. (2008). A stepwise model for double-stranded RNA processing by ribonuclease III. Mol. Microbiol. 67, 143–154.

Gleghorn, M. L., and Maquat, L. E. (2014). ‘Black sheep’ that don’t leave the double-stranded RNA-binding domain fold. Trends Biochem. Sci. 39, 328–340.

Gregory, R. I., Yan, K. P., Amuthan, G., Chendrimada, T., Doratotaj, B., Cooch, N., and Shiekhattar, R. (2004). The Microprocessor complex mediates the genesis of microRNAs. Nature 432, 235–240.

Grishok, A., Pasquinelli, A. E., Conte, D., Li, N., Parrish, S., Ha, I., Baillie, D. L., Fire, A., Ruvkun, G., and Mello, C. C. (2001). Genes and Mechanisms Related to RNA Interference Regulate Expression of the Small Temporal RNAs that Control *C. elegans* Developmental Timing. Cell 106, 23–34.

Gu, S., Jin, L., Zhang, Y., Huang, Y., Zhang, F., Valdmanis, P. N., and Kay, M. A. (2012). The loop position of shRNAs and pre-miRNAs is critical for the accuracy of dicer processing in vivo. Cell 151, 900–911.

Gurtan, A. M., Lu, V., Bhutkar, A., and Sharp, P. A. (2012). In vivo structure-function analysis of human Dicer reveals directional processing of precursor miRNAs. RNA 18, 1116–1122.

Haase, A. D., Jaskiewicz, L., Zhang, H., Lainé, S., Sack, R., Gatignol, A., and Filipowicz, W. (2005). TRBP, a regulator of cellular PKR and HIV-1 virus expression, interacts with Dicer and functions in RNA silencing. EMBO Rep. 6, 961–967.

Han, J., Lee, Y., Yeom, K. H., Kim, Y. K., Jin, H., and Kim, V. N. (2004). The Drosha- DGCR8 complex in primary microRNA processing. Genes Dev. 18, 3016–3027.

Han, J., Lee, Y., Yeom, K. H., Nam, J. W., Heo, I., Rhee, J. K., Sohn, S. Y., Cho, Y., Zhang, B. T., and Kim, V. N. (2006). Molecular basis for the recognition of primary microRNAs by the Drosha-DGCR8 complex. Cell 125, 887–901.

Hansen, S. R., Aderounmu, A. M., Donelick, H. M., and Bass, B. L. (2019). Dicer’s Helicase Domain: A Meeting Place for Regulatory Proteins. Cold Spring Harb. Symp. Quant. Biol. 84, 185–193.

Holbrook, S. R., Cheong, C., Tinoco, I., and Kim, S. H. (1991). Crystal structure of an RNA double helix incorporating a track of non-Watson-Crick base pairs. Nature 353, 579–581.

Hubbard, S. J., and Thornton, J. M. (1993). naccess. Computer Program, http://www.bioinf.manchester.ac.uk/naccess/

Hutvágner, G., McLachlan, J., Pasquinelli, A. E., Balint, É., Tuschl, T., and Zamore, P. D. (2001). A cellular function for the RNA-interference enzyme Dicer in the maturation of the *let-7* small temporal RNA. Science 293, 834–838.

Jakob, L., Treiber, T., Treiber, N., Gust, A., Kramm, K., Hansen, K., Stotz, M., Wankerl, L., Herzog, F., Hannus, S., Grohmann, D., and Meister, G. (2016). Structural and functional insights into the fly microRNA biogenesis factor Loquacious. RNA 22, 383–396.

Jiang, F., Ye, X., Liu, X., Fincher, L., McKearin, D., and Liu, Q. (2005). Dicer-1 and R3D1-L catalyze microRNA maturation in Drosophila. Genes Dev. 19, 1674–1679.

Jumper, J., Evans, R., Pritzel, A., Green, T., Figurnov, M., Ronneberger, O., Tunyasuvunakool, K., Bates, R., Žídek, A., Potapenko, A., Bridgland, A., Meyer, C., Kohl, S. A. A., Ballard, A. J., Cowie, A., Romera-Paredes, B., Nikolov, S., Jain, R., Adler, J., Back, T., Petersen, S., Reiman, D., Clancy, E., Zielinski, M., Steinegger, M., Pacholska, M., Berghammer, T., Bodenstein, S., Silver, D., Vinyals, O., Senior, A. W., Kavukcuoglu, K., Kohli, P., and Hassabis, D. (2021). Highly accurate protein structure prediction with AlphaFold. Nature 596, 583–589.

Kawamata, T., Seitz, H., and Tomari, Y. (2009). Structural determinants of miRNAs for RISC loading and slicer-independent unwinding. Nat. Struct. Mol. Biol. 16, 953–960.

Khvorova, A., Reynolds, A., and Jayasena, S. D. (2003). Functional siRNAs and miRNAs exhibit strand bias. Cell 115, 209–216.

Knight, S. W., and Bass, B. L. (2001). A role for the RNase III enzyme DCR-1 in RNA interference and germ line development in *Caenorhabditis elegans*. Science 293, 2269–2271.

Kremer, J. R., Mastronarde, D. N., and McIntosh, J. R. (1996). Computer visualization of three-dimensional image data using IMOD. J. Struct. Biol. 116, 71–76.

Kurzynska-Kokorniak, A., Pokornowska, M., Koralewska, N., Hoffmann, W., Bienkowska-Szewczyk, K., and Figlerowicz, M. (2016). Revealing a new activity of the human Dicer DUF283 domain in vitro. Scientific Reports 6, 23989.

Kwon, S., Nguyen, T., Choi, Y.-G., Jo, M., Hohng, S., Kim, V., and Woo, J.-S. (2016). Structure of Human DROSHA. Cell 164, 81–90.

Lai, E. C. (2002). Micro RNAs are complementary to 3’ UTR sequence motifs that mediate negative post-transcriptional regulation. Nat. Genet. 30, 363–364.

Lau, P.-W., Potter, C. S., Carragher, B., and MacRae, I. J. (2009). Structure of the Human Dicer-TRBP Complex by Electron Microscopy. Structure 17, 1326–1332.

Lau, P. W., Guiley, K. Z., De, N., Potter, C. S., Carragher, B., and MacRae, I. J. (2012). The molecular architecture of human Dicer. Nat. Struct. Mol. Biol. 19, 436–440.

Lee, Y., Ahn, C., Han, J., Choi, H., Kim, J., Yim, J., Lee, J., Provost, P., Rådmark, O., Kim, S., and Kim, V. N. (2003). The nuclear RNase III Drosha initiates microRNA processing. Nature 425, 415–419.

Lee, Y., Hur, I., Park, S.-Y., Kim, Y.-K., Suh, M. R., and Kim, V. N. (2006). The role of PACT in the RNA silencing pathway. EMBO J. 25, 522–532.

Lewis, B. P., Shih, I. H., Jones-Rhoades, M. W., Bartel, D. P., and Burge, C. B. (2003). Prediction of mammalian microRNA targets. Cell 115, 787–798.

Lim, M. Y., Ng, A. W., Chou, Y., Lim, T. P., Simcox, A., Tucker-Kellogg, G., and Okamura, K. (2016). The Drosophila Dicer-1 Partner Loquacious Enhances miRNA Processing from Hairpins with Unstable Structures at the Dicing Site. Cell Rep 15, 1795–1808.

Lingel, A., Simon, B., Izaurralde, E., and Sattler, M. (2003). Structure and nucleic-acid binding of the Drosophila Argonaute 2 PAZ domain. Nature 426, 465–469.

Lingel, A., Simon, B., Izaurralde, E., and Sattler, M. (2004). Nucleic acid 3’-end recognition by the Argonaute2 PAZ domain. Nat. Struct. Mol. Biol. 11, 576–577.

Liu, Z., Wang, J., Cheng, H., Ke, X., Sun, L., Zhang, Q. C., and Wang, H. W. (2018). Cryo-EM Structure of Human Dicer and Its Complexes with a Pre-miRNA Substrate. Cell 173, 1191–1203.e12.

Lorenz, R., Bernhart, S. H., Höner Zu Siederdissen, C., Tafer, H., Flamm, C., Stadler, P. F., and Hofacker, I. L. (2011). ViennaRNA Package 2.0. Algorithms Mol Biol 6, 26.

Luo, Q. J., Zhang, J., Li, P., Wang, Q., Zhang, Y., Roy-Chaudhuri, B., Xu, J., Kay, M. A., and Zhang, Q. C. (2021). RNA structure probing reveals the structural basis of Dicer binding and cleavage. Nat Commun 12, 3397.

Ma, J. B., Ye, K., and Patel, D. J. (2004). Structural basis for overhang-specific small interfering RNA recognition by the PAZ domain. Nature 429, 318–322.

MacRae, I. J., Zhou, K., and Doudna, J. A. (2007). Structural determinants of RNA recognition and cleavage by Dicer. Nat. Struct. Mol. Biol. 14, 934–940.

Macrae, I. J., Zhou, K., Li, F., Repic, A., Brooks, A. N., Cande, W. Z., Adams, P. D., and Doudna, J. A. (2006). Structural basis for double-stranded RNA processing by Dicer. Science 311, 195–198.

Maniataki, E., and Mourelatos, Z. (2005). A human, ATP-independent, RISC assembly machine fueled by pre-miRNA. Genes Dev. 19, 2979–2990.

Mastronarde, D. N. (2005). Automated electron microscope tomography using robust prediction of specimen movements. J. Struct. Biol. 152, 36–51.

Meister, G., Landthaler, M., Patkaniowska, A., Dorsett, Y., Teng, G., and Tuschl, T. (2004). Human Argonaute2 mediates RNA cleavage targeted by miRNAs and siRNAs. Mol. Cell. 15, 185–197.

Meng, W., and Nicholson, A. W. (2008). Heterodimer-based analysis of subunit and domain contributions to double-stranded RNA processing by Escherichia coli RNase III in vitro. Biochem. J. 410, 39–48.

Nicholson, A. W. (2014). Ribonuclease III mechanisms of double-stranded RNA cleavage. Wiley Interdiscip Rev RNA 5, 31–48.

Okamura, K., Ishizuka, A., Siomi, H., and Siomi, M. C. (2004). Distinct roles for Argonaute proteins in small RNA-directed RNA cleavage pathways. Genes Dev. 18, 1655–1666.

Ota, H., Sakurai, M., Gupta, R., Valente, L., Wulff, B.-E., Ariyoshi, K., Iizasa, H., Davuluri, R., and Nishikura, K. (2013). ADAR1 Forms a Complex with Dicer to Promote MicroRNA Processing and RNA-Induced Gene Silencing. Cell 153, 575–589.

Park, J.-E., Heo, I., Tian, Y., Simanshu, D. K., Chang, H., Jee, D., Patel, D. J., and Kim, V. N. (2011). Dicer recognizes the 5′ end of RNA for efficient and accurate processing. Nature 475, 201–205.

Pettersen, E. F., Goddard, T. D., Huang, C. C., Meng, E. C., Couch, G. S., Croll, T. I., Morris, J. H., and Ferrin, T. E. (2021). UCSF ChimeraX: Structure visualization for researchers, educators, and developers. Protein Sci 30, 70–82.

Provost, P. (2002). Ribonuclease activity and RNA binding of recombinant human Dicer. EMBO J. 21, 5864–5874.

Punjani, A., Zhang, H., and Fleet, D. J. (2020). Non-uniform refinement: adaptive regularization improves single-particle cryo-EM reconstruction. Nat Methods 17, 1214–1221.

Qin, H., Chen, F., Huan, X., Machida, S., Song, J., and Yuan, Y. A. (2010). Structure of the Arabidopsis thaliana DCL4 DUF283 domain reveals a noncanonical double-stranded RNA-binding fold for protein-protein interaction. RNA 16, 474–481.

Rohou, A., and Grigorieff, N. (2015). CTFFIND4: Fast and accurate defocus estimation from electron micrographs. J. Struct. Biol. 192, 216–221.

Saito, K., Ishizuka, A., Siomi, H., and Siomi, M. C. (2005). Processing of pre-microRNAs by the Dicer-1-Loquacious complex in Drosophila cells. PLoS Biol. 3, e235.

Savir, Y., and Tlusty, T. (2007). Conformational proofreading: the impact of conformational changes on the specificity of molecular recognition. PLoS One 2, e468.

Schwarz, D. S., Hutvágner, G., Du, T., Xu, Z., Aronin, N., and Zamore, P. D. (2003). Asymmetry in the assembly of the RNAi enzyme complex. Cell 115, 199–208.

Sheng, J., Larsen, A., Heuberger, B. D., Blain, J. C., and Szostak, J. W. (2014). Crystal structure studies of RNA duplexes containing s(2)U:A and s(2)U:U base pairs. J. Am. Chem. Soc. 136, 13916–13924.

Sinha, N. K., Iwasa, J., Shen, P. S., and Bass, B. L. (2018). Dicer uses distinct modules for recognizing dsRNA termini. Science 359, 329–334.

Song, J. J., Liu, J., Tolia, N. H., Schneiderman, J., Smith, S. K., Martienssen, R. A., Hannon, G. J., and Joshua-Tor, L. (2003). The crystal structure of the Argonaute2 PAZ domain reveals an RNA binding motif in RNAi effector complexes. Nat. Struct. Biol. 10, 1026–1032.

Sun, W., Pertzev, A., and Nicholson, A. W. (2005). Catalytic mechanism of Escherichia coli ribonuclease III: kinetic and inhibitor evidence for the involvement of two magnesium ions in RNA phosphodiester hydrolysis. Nucleic Acids Res. 33, 807–815.

Taylor, D. W., Ma, E., Shigematsu, H., Cianfrocco, M. A., Noland, C. L., Nagayama, K., Nogales, E., Doudna, J. A., and Wang, H. W. (2013). Substrate-specific structural rearrangements of human Dicer. Nat. Struct. Mol. Biol. 20, 662–670.

Tian, Y., Simanshu, D. K., Ma, J. B., Park, J. E., Heo, I., Kim, V. N., and Patel, D. J. (2014). A phosphate-binding pocket within the platform-PAZ-connector helix cassette of human Dicer. Mol. Cell. 53, 606–616.

Tomari, Y., Du, T., and Zamore, P. D. (2007). Sorting of Drosophila small silencing RNAs. Cell 130, 299–308.

Tsutsumi, A., Kawamata, T., Izumi, N., Seitz, H., and Tomari, Y. (2011). Recognition of the pre-miRNA structure by Drosophila Dicer-1. Nat. Struct. Mol. Biol. 18, 1153–1158.

Wagner, T., Merino, F., Stabrin, M., Moriya, T., Antoni, C., Apelbaum, A., Hagel, P., Sitsel, O., Raisch, T., Prumbaum, D., Quentin, D., Roderer, D., Tacke, S., Siebolds, B., Schubert, E., Shaikh, T. R., Lill, P., Gatsogiannis, C., and Raunser, S. (2019). SPHIRE-crYOLO is a fast and accurate fully automated particle picker for cryo-EM. Commun Biol 2, 218.

Wagner, T., and Raunser, S. (2020). The evolution of SPHIRE-crYOLO particle picking and its application in automated cryo-EM processing workflows. Commun Biol 3, 61.

Wang, H.-W., Noland, C., Siridechadilok, B., Taylor, D. W., Ma, E., Felderer, K., Doudna, J. A., and Nogales, E. (2009). Structural insights into RNA processing by the human RISC-loading complex. Nature Structural & Molecular Biology 16, 1148–1153.

Wang, Q., Xue, Y., Zhang, L., Zhong, Z., Feng, S., Wang, C., Xiao, L., Yang, Z., Harris, C. J., Wu, Z., Zhai, J., Yang, M., Li, S., Jacobsen, S. E., and Du, J. (2021). Mechanism of siRNA production by a plant Dicer-RNA complex in dicing-competent conformation. Science 374, 1152–1157.

Wei, X., Ke, H., Wen, A., Gao, B., Shi, J., and Feng, Y. (2021). Structural basis of microRNA processing by Dicer-like 1. Nat Plants 7, 1389–1396.

Wilson, R. C., Tambe, A., Kidwell, M. A., Noland, C. L., Schneider, C. P., and Doudna, J. A. (2015). Dicer-TRBP complex formation ensures accurate mammalian microRNA biogenesis. Mol. Cell. 57, 397–407.

Yan, K. S., Yan, S., Farooq, A., Han, A., Zeng, L., and Zhou, M. M. (2003). Structure and conserved RNA binding of the PAZ domain. Nature 426, 468–474.

Yang, S. W., Chen, H. Y., Yang, J., Machida, S., Chua, N. H., and Yuan, Y. A. (2010). Structure of Arabidopsis HYPONASTIC LEAVES1 and its molecular implications for miRNA processing. Structure 18, 594–605.

Ye, X., Paroo, Z., and Liu, Q. (2007). Functional anatomy of the Drosophila microRNA- generating enzyme. J. Biol. Chem. 282, 28373–28378.

Zapletal, D., Taborska, E., Pasulka, J., Malik, R., Kubicek, K., Zanova, M., Much, C., Sebesta, M., Buccheri, V., Horvat, F., Jenickova, I., Prochazkova, M., Prochazka, J., Pinkas, M., Novacek, J., Joseph, D. F., Sedlacek, R., O’Carroll, D., Stefl, R., and Svoboda, P. (2022). Molecular basis of indispensable accuracy of mammalian miRNA biogenesis. bioRxiv https://doi.org/10.1101/2022.04.13.488181.

Zhang, H., Kolb, F. A., Jaskiewicz, L., Westhof, E., and Filipowicz, W. (2004). Single processing center models for human Dicer and bacterial RNase III. Cell 118, 57–68.

Zhang, L., Xiang, Y., Chen, S., Shi, M., Jiang, X., He, Z., and Gao, S. (2022). Mechanisms of MicroRNA Biogenesis and Stability Control in Plants. Front Plant Sci 13, 844149.

Zheng, S. Q., Palovcak, E., Armache, J. P., Verba, K. A., Cheng, Y., and Agard, D. A. (2017). MotionCor2: anisotropic correction of beam-induced motion for improved cryo-electron microscopy. Nat Methods 14, 331–332.

Zhu, L., Kandasamy, S. K., and Fukunaga, R. (2018). Dicer partner protein tunes the length of miRNAs using base-mismatch in the pre-miRNA stem. Nucleic Acids Res. 46, 3726–3741.

Zivanov, J., Nakane, T., Forsberg, B. O., Kimanius, D., Hagen, W. J., Lindahl, E., and Scheres, S. H. (2018). New tools for automated high-resolution cryo-EM structure determination in RELION-3. eLife 7, e42166.

